# Structure and mechanism of the human CTDNEP1-NEP1R1 membrane protein phosphatase complex necessary to maintain ER membrane morphology

**DOI:** 10.1101/2023.11.20.567952

**Authors:** Shujuan Gao, Jake W. Carrasquillo Rodríguez, Shirin Bahmanyar, Michael V. Airola

**Author notes:** Address correspondence to: Michael V. Airola.

## Abstract

C-terminal Domain Nuclear Envelope Phosphatase 1 (CTDNEP1) is a non-canonical protein serine/threonine phosphatase that regulates ER membrane biogenesis. Inactivating mutations in CTDNEP1 correlate with development of medulloblastoma, an aggressive childhood cancer. The transmembrane protein Nuclear Envelope Phosphatase 1 Regulatory Subunit 1 (NEP1R1) binds CTDNEP1, but the molecular details by which NEP1R1 regulates CTDNEP1 function are unclear. Here, we find that knockdown of CTDNEP1 or NEP1R1 in human cells generate identical phenotypes, establishing CTDNEP1-NEP1R1 as an evolutionarily conserved membrane protein phosphatase complex that restricts ER expansion. Mechanistically, NEP1R1 acts as an activating regulatory subunit that directly binds and increases the phosphatase activity of CTDNEP1. By defining a minimal NEP1R1 domain sufficient to activate CTDNEP1, we determine high resolution crystal structures of the CTDNEP1-NEP1R1 complex bound to a pseudo-substrate. Structurally, NEP1R1 engages CTDNEP1 at a site distant from the active site to stabilize and allosterically activate CTDNEP1. Substrate recognition is facilitated by a conserved Arg residue that binds and orients the substrate peptide in the active site. Together, this reveals mechanisms for how NEP1R1 regulates CTDNEP1 and explains how cancer-associated mutations inactivate CTDNEP1.

## INTRODUCTION

A key branching point in de novo phospholipid synthesis is the conversion of phosphatidic acid (PA) to diacylglycerol by lipin PA phosphatases (PAPs), which promotes synthesis of the major membrane phospholipids and triglycerides^1,2^. The phosphorylation state of lipin regulates both PAP activity and subcellular localization^3–6^, with phosphorylation/dephosphorylation establishing a conserved regulatory mechanism between mammalian lipin PAPs and the orthologous yeast PAP *S. cerevisiae (Sc)* Pah1^7–9^. Several kinases, including mTOR can phosphorylate lipin^3,4,10,11^. C-terminal Domain Nuclear Envelope Phosphatase 1 (CTDNEP1) is the primary phosphatase that dephosphorylates lipin to regulate lipid and membrane biosynthesis^12–14^.

CTDNEP1 localizes to endoplasmic reticulum/nuclear envelope (ER/NE) membranes^15,16^ and plays key roles in regulating ER/NE membrane biogenesis^11,12^, nuclear pore insertion^17^, nuclear positioning^18,19^, and chromosome segregation during mitosis^11,20^. Recently, CTDNEP1 has been identified as a tumor suppressor with loss of function mutations in CTDNEP1 leading to amplification of c-MYC associated with medulloblastoma^20,21^, an aggressive childhood brain cancer.

CTDNEP1 is a member of the C-terminal domain phosphatases (CTDPs)^22^, which is a subfamily of the haloacid dehalogenase (HAD) superfamily of magnesium dependent phosphatases that share a characteristic Rossmann-like fold and catalytic mechanism involving a DxDx(V ⁄T) active site motif^12,14,23,24^. CTDNEP1 and CTDPs represent a distinct class of phosphoprotein phosphatases (PPPs) from canonical Ser/Thr PPPs (PP1-PP7) with different catalytic cores, metal ion requirements, and reaction mechanisms^22,25^. However, like canonical phosphatases, CTDNEP1 has been demonstrated to interact with other proteins^13,18^, which may act as regulatory subunits. Nuclear Envelope Phosphatase 1 Regulatory Subunit 1 (NEP1R1) is currently the best characterized binding partner of CTDNEP1^13,15–17^.

The yeast orthologs of CTDNEP1 and NEP1R1 in *S. cerevisiae* are Nem1 and Spo7, which form an obligatory phosphatase complex that dephosphorylates Pah1^9,26–28^, the yeast ortholog of lipins. Genetic disruption of *NEM1* or *SPO7* give rise to identical phenotypes with an expanded nuclear envelope morphology and defects in sporulation^26^. Co-expression of NEP1R1 with CTDNEP1 is required to rescue these effects in *spo7*Δ or *nem1*Δ *spo7*Δ cells^13^, but expression of CTDNEP1 alone is sufficient to complement *nem1*Δ cells^12^. In mammalian cells, the requirement of NEP1R1 co-expression for CTDNEP1 to dephosphorylate lipin PAPs is cell type dependent, with NEP1R1 required in HEK293 cells, but not in BHK cells^12^. While it is clear NEP1R1 can function as a partner for CTDNEP1 to dephosphorylate lipin, the mechanistic role of NEP1R1 remains undefined.

Here, we address the mechanistic and structural role of NEP1R1 as a regulatory subunit for CTDNEP1. We find NEP1R1, like CTDNEP1^11^, is also required to limit ER expansion in mammalian cells. Extensive biochemical data, using highly purified proteins, defines NEP1R1 as an activating transmembrane regulatory subunit that associates with CTDNEP1 through a soluble, non-membrane embedded domain with micromolar affinity. High resolution crystal structures of the CTDNEP1-NEP1R1 complex, with an active site bound pseudo-substrate peptide, reveal the structural basis of complex formation and substrate recognition, and explain how cancer associated mutations inactivate the protein phosphatase activity of human CTDNEP1.

## RESULTS

### Interdependency of CTDNEP1 and NEP1R1 function in mammalian cells

To address the role of NEP1R1, we first generated a U2OS CTDNEP1 KO cell line stably expressing epitope-tagged CTDNEP1 and NEP1R1 and used siRNA to monitor the co-dependence of protein levels. As previously observed upon overexpression in yeast or HEK293 cells^13^, CTDNEP1 and NEP1R1 protein levels were interdependent, with knockdown of either protein reducing the levels of both proteins **(Fig. 1A)**.

**Fig. 1.**
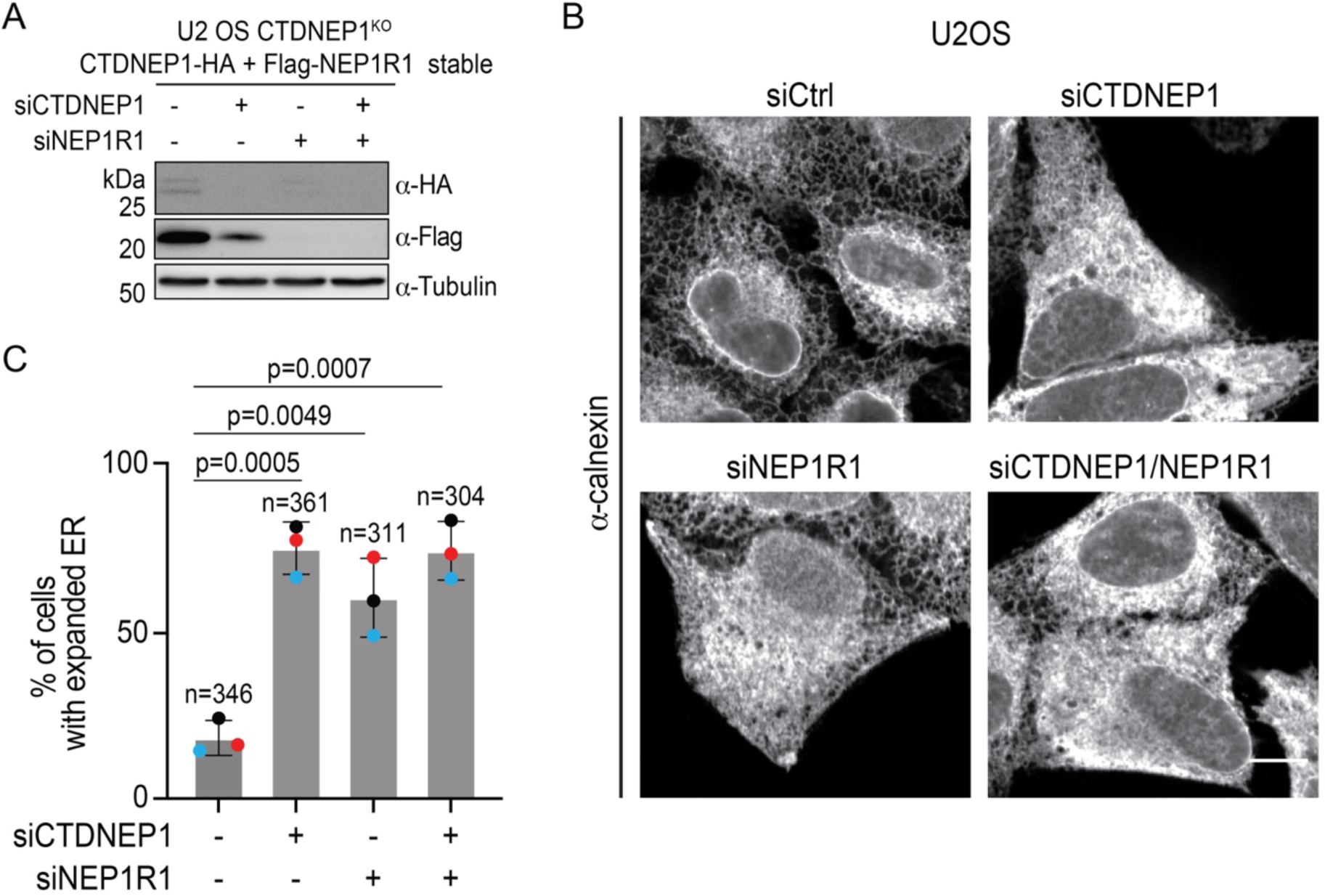
Interdependency of CTDNEP1 and NEP1R1 protein stability and function in U2OS cells. **(A)** Immunoblot of whole cell lysates from U2OS CTDNEP1^KO^ cells stably expressing CTDNEP1-HA and Flag-NEP1R1, siRNA treated as indicated, (N = 2 independent experiments). **(B)** Representative spinning disk confocal images of U2OS cells under the indicated siRNA conditions, immunostained with anti-calnexin. scale bars, 10 μm. **(C)** Plot, % of cells with expanded ER under indicated siRNA conditions from Figure B. Bars indicate mean ± SDs (N = 3, independent experimental repeats as shown by colored dots, n indicates the number of cells quantified). p-values were determined by an unpaired t-test.

To assess the functional consequences of NEP1R1 knockdown, we took advantage of our recent observation that knockout of CTDNEP1 results in an expanded ER phenotype in U2OS cells^11^. As seen with the CTDNEP1 KO^11^, control U2OS cells treated with CTDNEP1 siRNA had an expanded ER in comparison to non-targeting control siRNA **(Fig. 1B, C)**. Similarly, siRNA depletion of NEP1R1 significantly elevated the percentage of cells with expanded ER **(Fig. 1B, C)**. Knockdown of both CTDNEP1 and NEP1R1 did not result in a further increase of ER expansion **(Fig. 1B, C)**. This indicates a functional interdependence of CTDNEP1 and NEP1R1 in limiting ER expansion.

### CTDNEP1 and NEP1R1 form a catalytically active magnesium-dependent protein phosphatase complex

Given their functional interdependence in limiting ER expansion, we sought to determine the mechanism by which NEP1R1 acts as a regulatory subunit for CTDNEP1. The role of NEP1R1 has been unclear given CTDNEP1 is catalytically active in vitro in the absence of NEP1R1^12,14^ and the difficulties associated with purification of the transmembrane protein NEP1R1.

We first purified and compared the activity of the NEP1R1/CTDNEP1 protein complex with CTDNEP1 alone. Highly pure NEP1R1/CTDNEP1 complex was obtained by co-expression of a His-tagged maltose binding protein (MBP) fusion of NEP1R1 with untagged CTDNEP1 (MBP-NEP1R1/CTDNEP1) in *E. coli*, followed by purification using Ni-NTA and size-exclusion chromatography **(Fig. 2A, B)**. As a control, we purified an inactive point mutant of the complex that replaced the catalytic aspartate residue D67 of CTDNEP1 with glutamate (MBP-NEP1R1/CTDNEP1^D67E^) **(Fig. 2A, B)**. In both cases, untagged wild type (WT) CTDNEP1 and the D67E point mutant remained bound to NEP1R1 throughout the purification process.

**Fig. 2.**
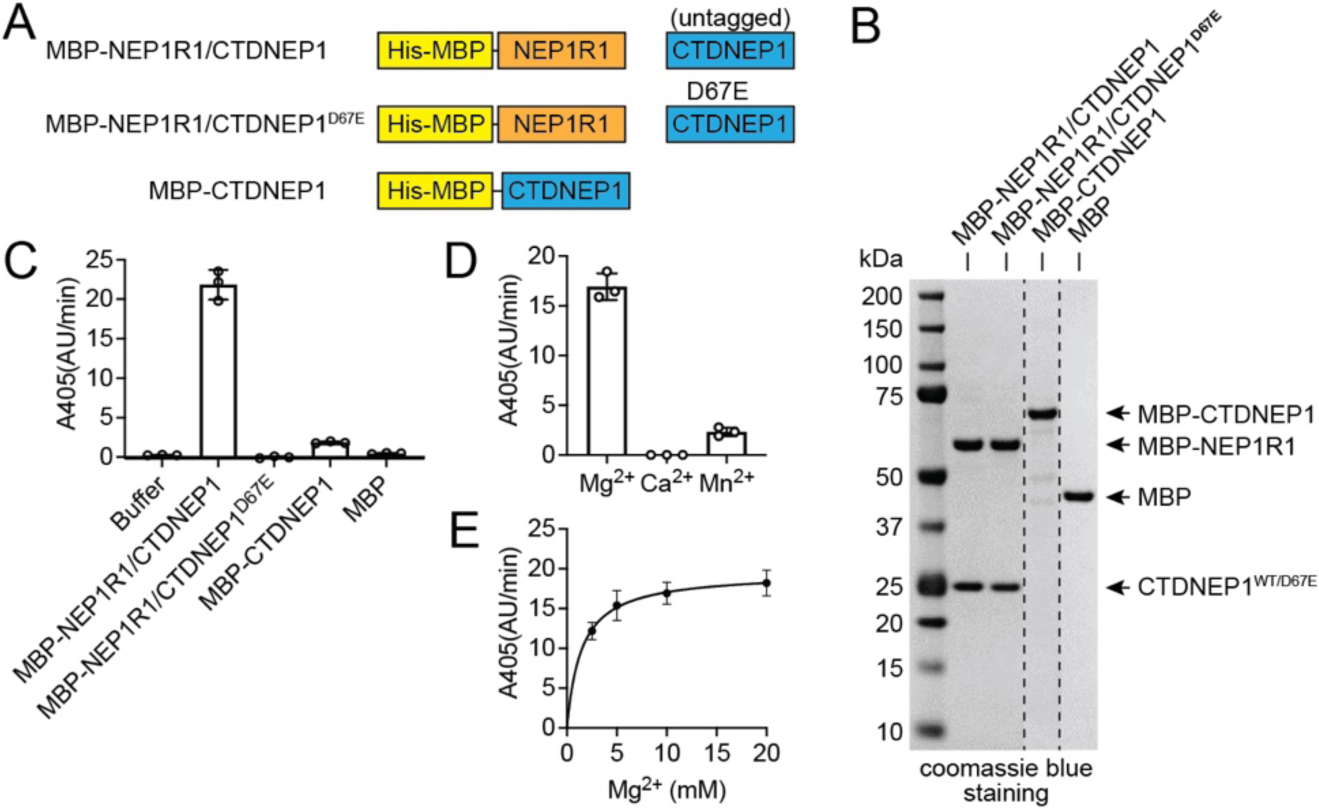
NEP1R1 forms an active phosphatase complex with CTDNEP1. **(A)** Domain architecture of purified CTDNEP1 and NEP1R1 proteins. MBP, maltose binding protein; His, His-tag. CTDNEP1 required fusion with MBP for purification unless co-expressed with NEP1R1. **(B)** SDS-PAGE analysis of purified CTDNEP1 and NEP1R1 proteins using Coomassie blue stain. **(C)** Quantification of pNPP hydrolysis by CTDNEP1 alone or in complex with NEP1R1. The D67E active site point mutant eliminated activity. Error bars represent standard deviation (*n* = 3). **(D)** Effect of metal ions (10 mM) on MBP-NEP1R1/CTDNEP1 complex activity using pNPP as a substrate. Error bars represent standard deviation (*n* = 3). **(E)** The activity of the MBP-NEP1R1/CTDNEP1 complex towards pNPP depends on magnesium concentration. Error bars represent standard deviation (*n* = 3).

On its own, purification of full-length CTDNEP1 required fusion with MBP (MBP-CTDNEP1). Cleavage of MBP caused the resulting untagged CTDNEP1 to precipitate. We thus limited our characterization to MBP-CTDNEP1. When purified alone, MBP-CTDNEP1 formed a soluble aggregate on size-exclusion chromatography, while the co-purified MBP-NEP1R1/CTDNEP1 protein complex did not aggregate. This indicated that association with NEP1R1 prevents aggregation of CTDNEP1 in vitro.

The MBP-NEP1R1/CTDNEP1 complex was catalytically active and robustly hydrolyzed the generic phosphatase substrate para-nitrophenol phosphate (pNPP) **(Fig. 2C)**. pNPP hydrolysis by MBP-NEP1R1/CTDNEP1 was magnesium dependent with manganese generating some activity and calcium failing to support any catalytic function **(Fig. 2D, E)**. MBP-CTDNEP1 was capable of hydrolyzing pNPP but had ∼10-fold lower activity than the MBP-NEP1R1/CTDNEP1 complex. As expected, the negative controls MBP-NEP1R1/CTDNEP1^D67E^ and MBP did not hydrolyze pNPP **(Fig. 2C)**. We concluded that NEP1R1 and CTDNEP1 form an active magnesium-dependent phosphatase complex, and that in vitro NEP1R1 stabilizes and prevents aggregation of CTDNEP1.

### CTDNEP1 binds membranes via an N-terminal amphipathic helix

CTDNEP1 belongs to the HAD superfamily of phosphatase enzymes^12,14^. Alphafold^29^ predicts CTDNEP1 to form a globular HAD-like catalytic domain with an N-terminal amphipathic helix that contains hydrophobic residues lining one side of the helix and polar residues on the other side **(Fig. 3A, B).** This suggests CTDNEP1 membrane association is mediated by the predicted N-terminal amphipathic helix (AH), which has previously been proposed to be a transmembrane helix^12,14^.

**Fig. 3.**
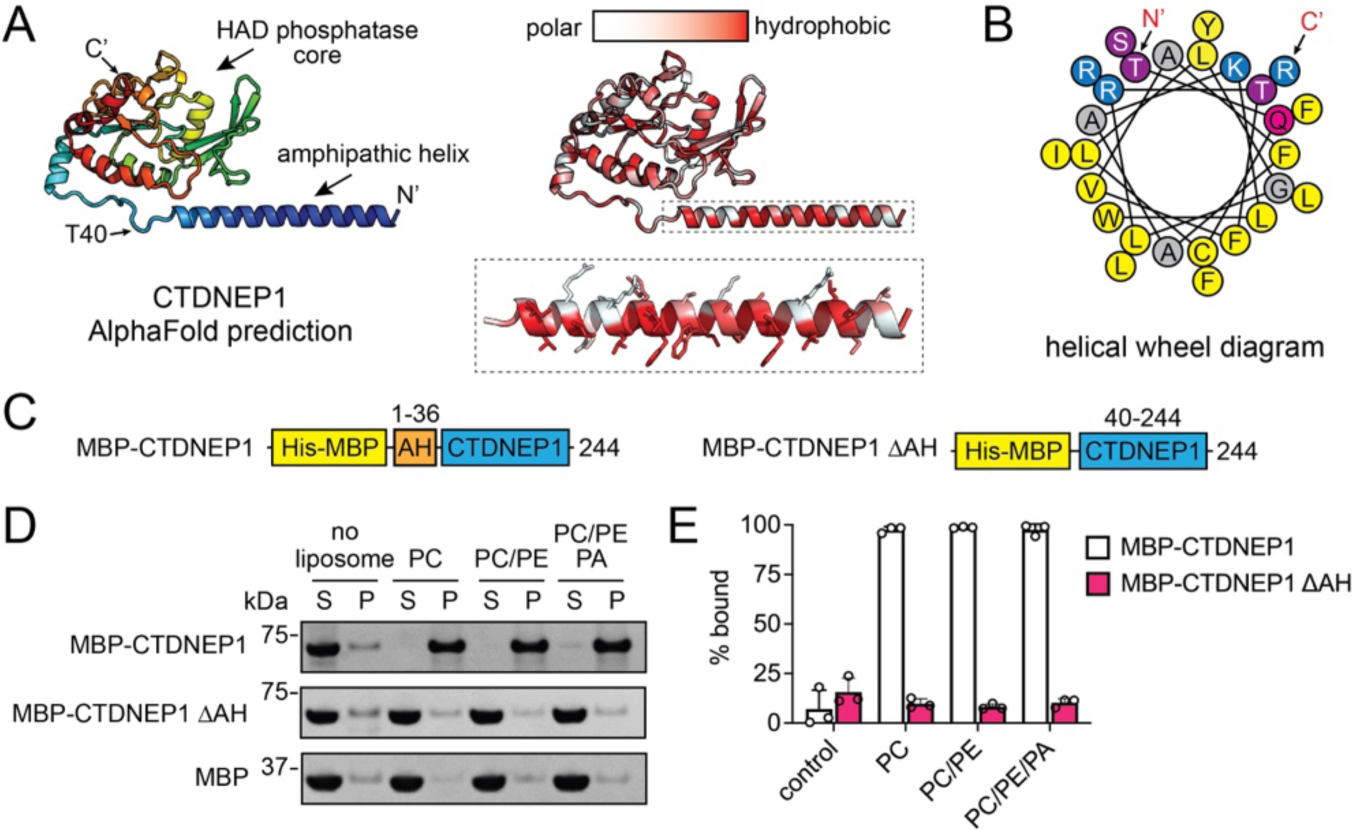
An N-terminal amphipathic helix mediates CTDNEP1 membrane recruitment. **(A)** Alphafold2 predicts human CTDNEP1 to form a globular catalytic domain from the HAD phosphatase family with an N-terminal amphipathic helix. The predicted CTDNEP1 structure is shown in cartoon form with rainbow coloring (left) from the N-terminal (blue) to C-terminal (red), or by hydrophobicity (right) with hydrophobic residues in red and hydrophilic residues in white. The inset depicts the sidechains of the N-terminal amphipathic helix. **(B)** Helical wheel diagram of the CTDNEP1 N-terminal amphipathic helix with bulky and small hydrophobic residues in yellow and grey, polar residues in purple or magenta, and positively charged residues in blue. The N- and C-termini positions are indicated. **(C)** Domain architecture of purified MBP-CTDNEP1 fusion proteins with and without the N-terminal amphipathic helix. **(D)** SDS-PAGE analysis of liposome co-sedimentation assays reveals CTDNEP1 strongly associates with membranes irrespective of lipid composition. Deletion of the N-terminal amphipathic helix in CTDNEP1 (MBP-CTDNEP1ΔAH) results in complete loss of membrane association. S, supernatant; P, pellet; PC, phosphatidylcholine; PE, phosphatidylethanolamine; PA, phosphatidic acid; MBP, maltose binding protein. **(E)** Quantification of liposome association for MBP-CTDNEP1 wild-type and ΔAH. Error bars represent standard deviation (*n* = 3).

To test if the amphipathic helix was required for CTDNEP1 membrane association, we purified CTDNEP1 lacking this helix (MBP-CTDNEP1ΔAH) **(Fig. 3C)** and used liposome co-sedimentation to assess membrane binding. Under all lipid compositions tested, MBP-CTDNEP1 displayed ∼100% binding to liposomes and deletion of the amphipathic helix eliminated all membrane binding **(Fig. 3D, 3E)**. We concluded that CTDNEP1 contains an N-terminal amphipathic helix that is required for and mediates membrane association.

### NEP1R1 binds and enhances the phosphatase activity of CTDNEP1

The ability of CTDNEP1 to dephosphorylate lipin in cells has been shown to require NEP1R1^13^. However, two independent studies have demonstrated recombinant CTDNEP1 alone exhibits in vitro catalytic activity against pNPP^12^ or 9-mer phospho-peptides of lipin^14^. Both in vitro studies used recombinant CTDNEP1 that lacked the amphipathic helix^12,14^. We purified NEP1R1 separately to test if NEP1R1 could bind and activate CTDNEP1 in vitro. As a comparison, we included co-purified complexes of MBP-NEP1R1/CTDNEP1 and MBP-NEP1R1/CTDNEP1ΔAH, which respectively contained or lacked the N-terminal amphipathic helix **(Fig. 4A)**.

**Fig. 4.**
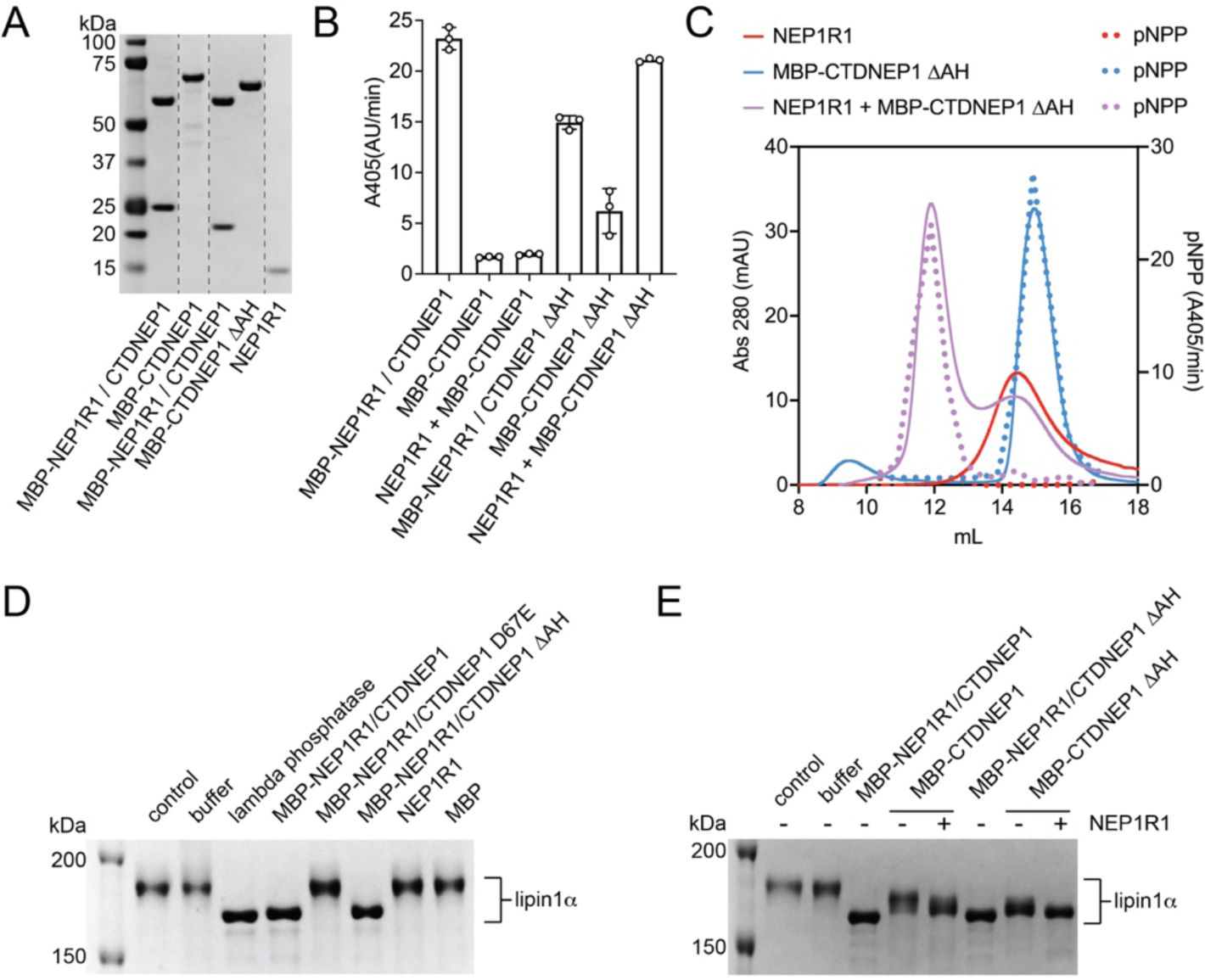
NEP1R1 binds and enhances the activity of CTDNEP1. **(A)** SDS-PAGE analysis of purified CTDNEP1 and NEP1R1 proteins stained with Coomassie blue. **(B)** Quantification of pNPP hydrolysis by MBP-CTDNEP1 alone, CTDNEP1 co-purified with NEP1R1 (MBP-NEP1R1/CTDNEP1), or with purified NEP1R1 added separately to MBP-CTDNEP1 (NEP1R1 + MBP-CTDNEP1 at 2:1 molar ratio). ΔAH indicates deletion of the CTDNEP1 N-terminal amphipathic helix. Error bars represent standard deviation (*n* = 3). **(C)** Size-exclusion profiles (solid lines) and quantification pNPP activity (dotted lines) of MBP-CTDNEP1ΔAH (blue traces) and NEP1R1 (red traces) alone, or a mixture of MBP-CTDNEP1ΔAH+NEP1R1 (purple traces) with NEP1R1 protein in 4x molar excess. The addition of NEP1R1 shifts both the elution profile and the fractions capable of hydrolyzing pNPP to a higher apparent molecular weight indicating complex formation between NEP1R1 and CTDNEP1. **(D)** Dephosphorylation by the NEP1R1-CTDNEP1 complex changes the migration of lipin 1α on phos-tag SDS-PAGE. **(E)** Phos-tag SDS-PAGE analysis of the effects of NEP1R1 on CTDNEP1-mediated dephosphorylation of lipin 1α.

The MBP-NEP1R1/CTDNEP1 complex had higher catalytic activity towards pNPP than the MBP-NEP1R1/CTDNEP1ΔAH complex **(Fig. 4B)**. However, NEP1R1 failed to increase activity of full length CTDNEP1 (MBP-CTDNEP1) **(Fig. 4B)**. We suspected this may be due to the observed aggregation of MBP-CTDNEP1 when purified in the absence of NEP1R1. Consistently, MBP-CTDNEP1ΔAH, which lacked the amphipathic helix and did not aggregate, displayed higher activity than full length MBP-CTDNEP1. Notably, MBP-CTDNEP1ΔAH was activated by NEP1R1 to levels comparable to the co-purified complexes **(Fig. 4B)**. We concluded that NEP1R1 directly activates CTDNEP1, and that complex formation and phosphatase activity does not require CTDNEP1’s amphipathic helix.

To confirm that activation of CTDNEP1 by NEP1R1 was from a direct protein-protein interaction, we used size-exclusion chromatography (SEC) to assess binding between NEP1R1 and CTDNEP1. Elution fractions were tested for pNPP hydrolysis as a readout for the presence of CTDNEP1. When ran separately, NEP1R1 and MBP-CTDNEP1ΔAH had similar retention times to each other **(Fig. 4C)**. The NEP1R1 SEC peak did not hydrolyze pNPP, while the MBP-CTDNEP1ΔAH peak directly overlapped with fractions that hydrolyzed pNPP **(Fig. 4C)**.

A mixture of MBP-CTDNEP1ΔAH with a 4-molar excess of NEP1R1 gave rise to two non-overlapping peaks. One peak (elution volume, V_E_ = 14.4 mL) overlapped with free NEP1R1. A second larger apparent molecular weight peak (V_E_ = 11.9 mL) also appeared, which represented the NEP1R1/MBP-CTDNEP1ΔAH complex as confirmed by SDS-PAGE analysis **(Supplementary Fig. 1)** and the direct overlap of these fractions with pNPP hydrolysis **(Fig. 4C)**. Thus, activation of CTDNEP1 by NEP1R1 occurs through a direct protein-protein interaction and the amphipathic helix of CTDNEP1 is not required for complex formation in vitro.

We next compared the ability of NEP1R1 to regulate CTDNEP1 dephosphorylation of lipin, as lipin represents an evolutionary conserved well characterized phosphoprotein substrate of CTDNEP1^11–14^. These experiments used purified mouse lipin 1α expressed in Sf9 insect cells and took advantage of the gel shift observed upon lipin dephosphorylation^3,6,30^. At the single protein concentration tested, we observed near complete dephosphorylation of lipin 1α by the MBP-NEP1R1/CTDNEP1 complex that was comparable to treatment with lamda phosphatase, with no observable dependence on CTDNEP1’s amphipathic helix **(Fig. 4D)**. For CTDNEP1 alone, deletion of the helix (MBP-CTDNEP1ΔAH) resulted in a slightly faster migration of lipin 1α on the phos-tag gel **(Fig. 4E)**, which suggests a higher catalytic rate of the non-aggregated MBP-CTDNEP1ΔAH over MBP-CTDNEP1. Consistent with our previous observations, the addition of NEP1R1 further enhanced lipin 1α dephosphorylation by CTDNEP1 **(Fig. 4E)**. Thus, NEP1R1 directly binds and enhances the catalytic activity of CTDNEP1 towards both the artificial substrate pNPP and the canonical phosphoprotein substrate lipin 1α.

### A soluble cytoplasmic domain of NEP1R1 is sufficient to bind and activate CTDNEP1

We next sought to define a minimal region of NEP1R1 sufficient to bind and active CTDNEP1. Alphafold predicted NEP1R1 to adopt an elongated structure containing a transmembrane helical region and a cytosolic domain **(Fig. 5A**). Given CTDNEP1 is a peripheral membrane-binding protein, we reasoned that the interaction between NEP1R1 and CTDNEP1 might be through the non-membrane embedded, cytoplasmic domain of NEP1R1. This hypothesis was assessed by generating a soluble version of NEP1R1 that deleted NEP1R1’s predicted transmembrane helices **(Fig. 5A, B)**. This soluble construct of NEP1R1, which did not require detergents for purification, is referred to as soluble NEP1R1 (sNEP1R1).

**Fig. 5.**
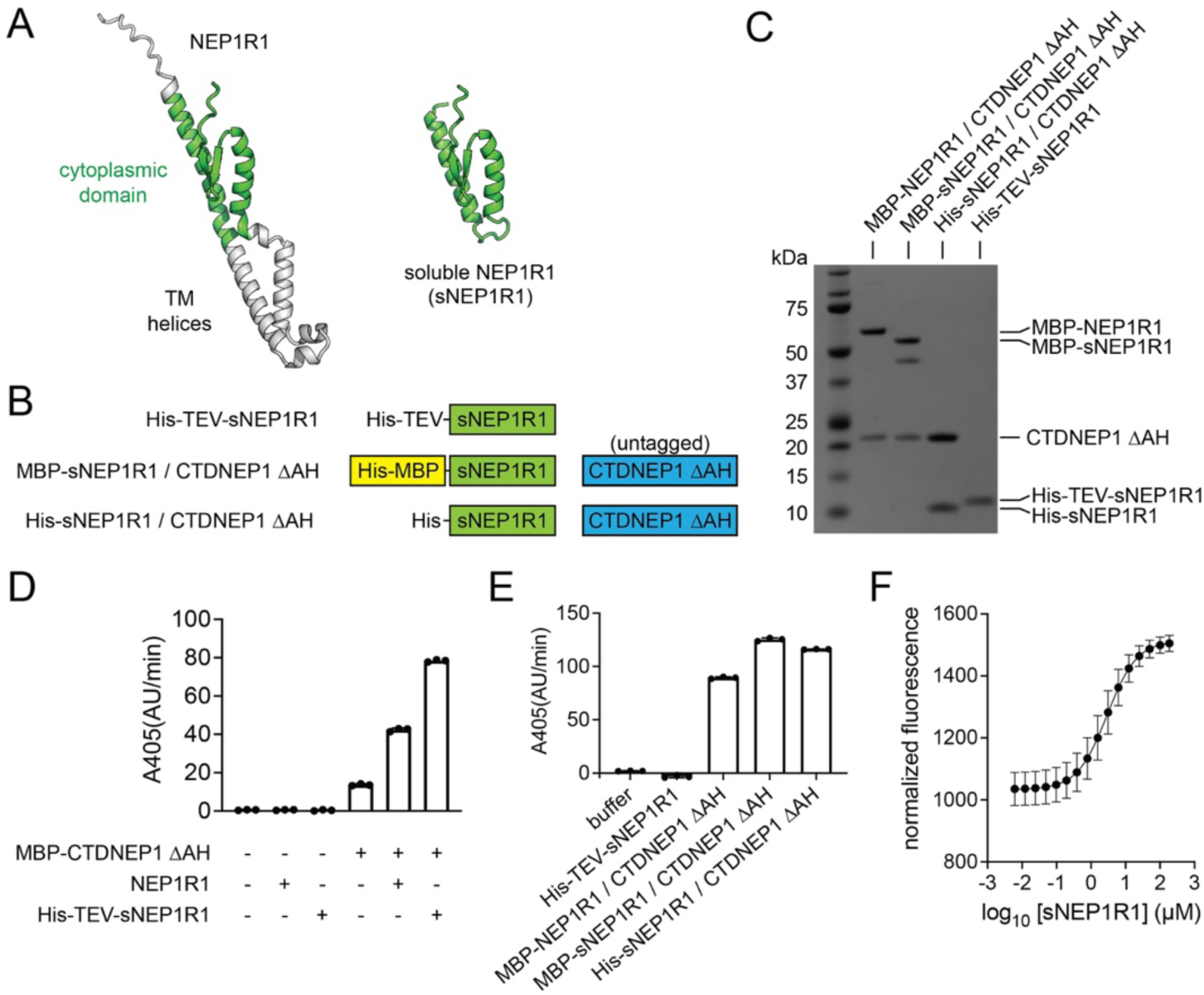
A soluble minimal cytoplasmic domain of NEP1R1 is sufficient to bind and activate CTDNEP1. **(A)** Cartoon representation of the predicted NEP1R1 structure with the cytoplasmic soluble domain in green. Removal of the TM helices and disordered N-terminus (grey) generates a soluble version of NEP1R1 (sNEP1R1). **(B)** Domain architectures of His-sNEP1R1, the MBP-sNEP1R1/CTDNEP1ΔAH complex, and the His-sNEP1R1/CTDNEP1ΔAH complex. **(C)** SDS-PAGE analysis of purified sNEP1R1 and the sNEP1R1/CTDNEP1ΔAH complexes using Coomassie blue stain. **(D)** Quantification of pNPP hydrolysis by MBP-CTDNEP1ΔAH alone, or with purified NEP1R1 (MBP-CTDNEP1ΔAH + NEP1R1 at 1:2 molar ratio) or sNEP1R1 (MBP-CTDNEP1ΔAH + His-TEV-sNEP1R1 at 1:2 molar ratio) added separately. Error bars represent standard deviation (*n* = 3). **(E)** Quantification of pNPP hydrolysis by CTDNEP1ΔAH co-purified with either NEP1R1 (MBP-NEP1R1/CTDNEP1ΔAH) or sNEP1R1 (MBP-sNEP1R1/CTDNEP1ΔAH or His-sNEP1R1/CTDNEP1ΔAH). Error bars represent standard deviation (*n* = 3). **(F)** Microscale thermophoresis indicates a dissociation constant (K_d_) of 2.9 μM between a msfGFP-fusion of MBP-CTDNEP1ΔAH and His-TEV-sNEP1R1. Error bars represent standard deviation (*n* = 4).

sNEP1R1 could be purified alone **(Fig. 5C)** and formed a stable domain that was sufficient to increase the activity of MBP-CTDNEP1ΔAH using pNPP as a substrate **(Fig. 5D)**. In addition, untagged CTDNEP1ΔAH co-purified with either MBP-sNEP1R1 or His-sNEP1R1 when co-expressed together in *E. coli* **(Fig. 5C)**. These soluble complexes were catalytically active **(Fig. 5E)** and band intensities suggested CTDNEP1 and sNEP1R1 associate in a 1 to 1 molar ratio **(Supplementary Fig. 2)**. This defines a minimal cytoplasmic domain of NEP1R1 that is sufficient to bind and activate CTDNEP1.

### NEP1R1 and CTDNEP1 associate with low micromolar affinity

To quantitate the affinity of the interaction between NEP1R1 and CTDNEP1, we used microscale thermophoresis (MST) to determine the equilibrium dissociation constant (K_d_). MST experiments were carried out by addition of increasing concentrations of sNEP1R1 to a solution of msfGFP-MBP-CTDNEP1ΔAH, a fusion of monomeric superfold green fluorescent protein (msfGFP) with MBP-CTDNEP1ΔAH. Fitting of the relative changes in thermophoresis yielded a K_d_ value of 2.9 μM **(Fig. 5F, Supplementary Fig. 3)**, which is in the physiological range of protein-protein interactions.

### Crystal structure of the CTDNEP1-NEP1R1 protein phosphatase complex

To determine the structural basis for CTDNEP1 activation by NEP1R1, we attempted to crystallize and/or use cryoEM to determine the structure of CTDNEP1 alone, as well as full-length and soluble CTDNEP1-NEP1R1 complexes, but were unsuccessful.

We thus generated several covalent fusions of CTDNEP1ΔAH with sNEP1R1 that contained different linker peptides. One of these fusions had similar activity as the soluble unfused complex **(Fig. 6A, B)** and crystallized, diffracting to 1.91 Å (R_work_ = 0.1908, R_free_ = 0.2120, **Table 1**). Phases were determined using molecular replacement with an Alphafold model of CTDNEP1, as molecular replaced failed with an Alphafold multimer^31^ model of the soluble complex. A high-resolution structure of the CTDNEP1-sNEP1R1 fusion was also determined with a magnesium (Mg^2+^) ion bound in the active site **(Table 1)**.

**Fig. 6.**
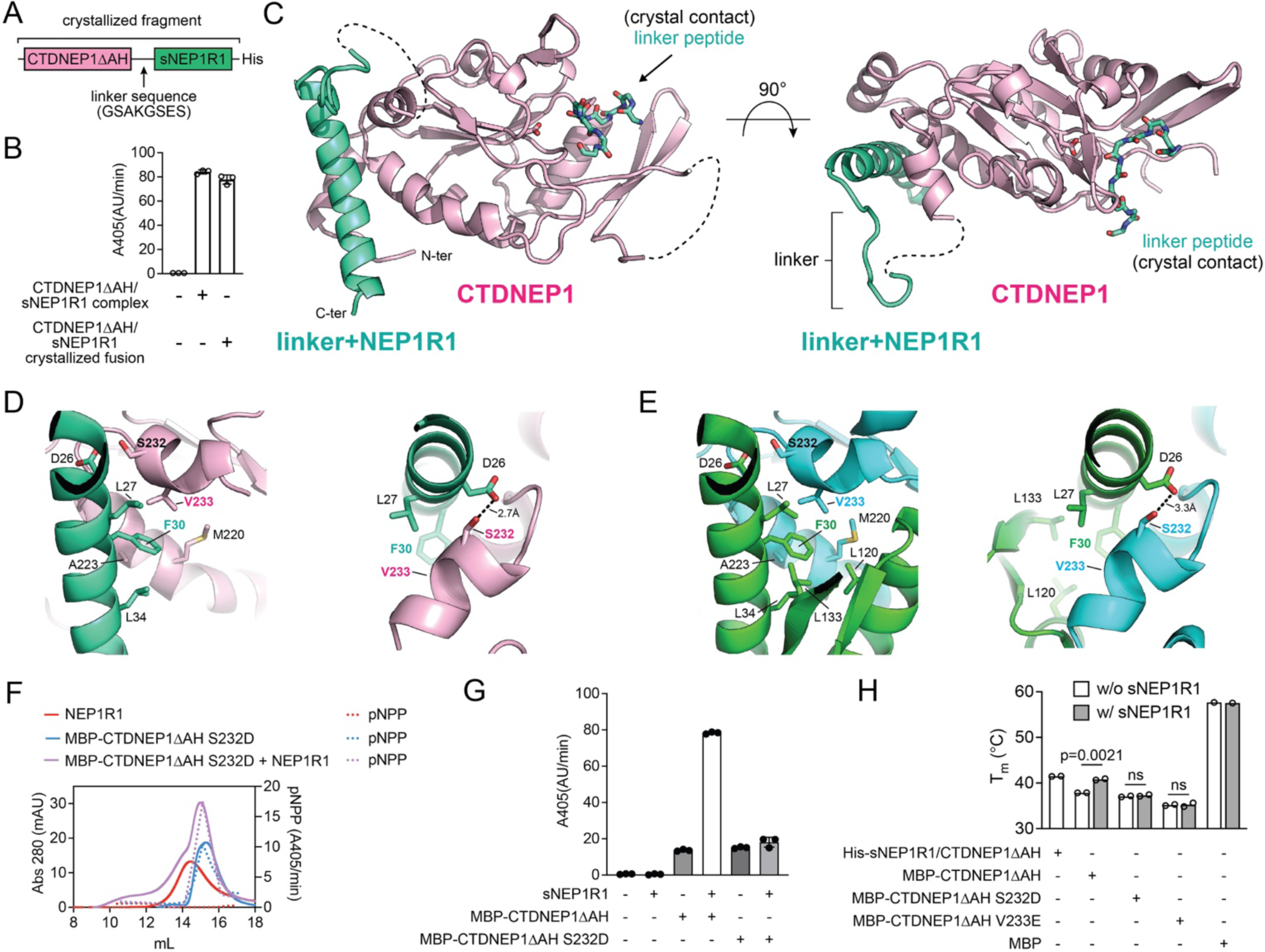
Crystal structure of the human CTDNEP1-NEP1R1 phosphatase complex. **(A)** Domain architecture of the crystallized CTDNEP1ΔAH-sNEP1R1 fusion with linker sequence. **(B)** Quantification of pNPP hydrolysis by CTDNEP1ΔAH /sNEP1R1 co-purified complex and the CTDNEP1ΔAH -sNEP1R1 fusion. Error bars represent standard deviation (*n* = 3). **(C)** Overall structure of the CTDNEP1-NEP1R1 phosphatase complex. A linker peptide occupies the active site of the CTDNEP1 catalytic subunit. NEP1R1 forms an extended helix that packs against the opposite side of the CTDNEP1 active site. **(D-E)** Interactions between NEP1R1 and CTDNEP1 **(D)** observed in the crystal structure and **(E)** predicted by AlphaFold multimer**. (F)** Size-exclusion profiles (solid lines) and quantification pNPP activity (dotted lines) of MBP-CTDNEP1ΔAHS232D (blue traces), NEP1R1 (red traces), and a mixture of CTDNEP1ΔAHS232D+NEP1R1 (purple traces). **(G)** Quantification of pNPP hydrolysis by MBP-CTDNEP1ΔAH WT and S232D with and without sNEP1R1. Error bars represent standard deviation (*n* = 3). **(H)** Melting temperatures (T_m_’s) of the His-sNEP1R1/CTDNEP1ΔAH complex in comparison to WT, S232D, and V233E MBP-CTDNEP1ΔAH with and without sNEP1R1. sNEP1R1 increases the stability of MBP-CTDNEP1ΔAH, but not the point mutants S232D and V233E that disrupt the complex. p-values were determined by an unpaired t-test. (n=2, or n=1 for MBP alone)

**Table 1.** Data collection and refinement statistics.

|  | CTDNEP1-NEP1R1 fusion | CTDNEP1-NEP1R1 fusion with Mg<sup>2+</sup> |
| --- | --- | --- |
| PDB code | 8UJL | 8UJM |
| Wavelength (Å) | 0.9201 | 0.9201 |
| Resolution range (Å) | 29.08 - 1.908 (1.976 - 1.908) | 50.45 - 2.161 (2.238 - 2.161) |
| Space group | C 2 2 21 | P 1 21 1 |
| Unit cell | 58.151 98.849 103.101 90 90 90 | 56.405 102.953 57.244 90 118.196 90 |
| Total reflections | 46893 (4520) | 61157 (5967) |
| Unique reflections | 23447 (2260) | 30755 (3010) |
| Multiplicity | 2.0 (2.0) | 2.0 (2.0) |
| Completeness (%) | 99.74 (98.01) | 99.54 (98.56) |
| Mean I/sigma(I) | 7.92 (0.83) | 5.67 (1.13) |
| Wilson B-factor | 32.92 | 33.65 |
| R-merge | 0.05073 (0.7953) | 0.09156 (0.7214) |
| R-meas | 0.07175 (1.125) | 0.1295 (1.02) |
| R-pim | 0.05073 (0.7953) | 0.09156 (0.7214) |
| CC1/2 | 0.998 (0.412) | 0.992 (0.385) |
| CC* | 1 (0.764) | 0.998 (0.746) |
| Reflections used in refinement | 23446 (2260) | 30744 (3010) |
| Reflections used for R-free | 1109 (128) | 1491 (161) |
| R-work | 0.1908 (0.3179) | 0.1969 (0.2868) |
| R-free | 0.2120 (0.3249) | 0.2205 (0.3180) |
| CC(work) | 0.954 (0.663) | 0.951 (0.659) |
| CC(free) | 0.958 (0.641) | 0.966 (0.653) |
| Number of non-hydrogen atoms | 1929 | 4054 |
| macromolecules | 1750 | 3777 |
| ligands | 0 | 2 |
| solvent | 179 | 275 |
| Protein residues | 218 | 472 |
| RMS(bonds) | 0.005 | 0.002 |
| RMS(angles) | 0.57 | 0.52 |
| Ramachandran favored (%) | 97.64 | 96.30 |
| Ramachandran allowed (%) | 1.89 | 3.48 |
| Ramachandran outliers (%) | 0.47 | 0.22 |
| Rotamer outliers (%) | 0.50 | 1.40 |
| Clashscore | 1.43 | 0.92 |
| Average B-factor | 44.79 | 47.13 |
| macromolecules | 44.29 | 47.27 |
| ligands | n/a | 35.23 |
| solvent | 49.71 | 45.22 |

Statistics for the highest-resolution shell are shown in parentheses.

The CTDNEP1-sNEP1R1 phosphatase complex had an overall compact structure, with CTDNEP1 adopting the expected globular HAD-phosphatase fold and sNEP1R1 forming a single extended helix that engaged CTDNEP1 on a hydrophobic surface present on the opposite side from the active site **(Fig. 6C)**. Notably, the hydrophobic surface in CTDNEP1 is not conserved in the similar human CTD phosphatases **(Supplementary Fig. 4)**, which are not known to bind regulatory subunits. This suggests the dependency of CTDNEP1 on NEP1R1 derives from this hydrophobic surface, which is occluded by NEP1R1 to stabilize and activate CTDNEP1.

The NEP1R1 helix engaging CTDNEP1 encompassed conserved region 1 (amino acids 24-38), which is one of three conserved regions between NEP1R1, Spo7, and other homologs based on secondary structure analysis^13^. However, several key residues at the interface of NEP1R1 and CTDNEP1 complex are not conserved in *S. cerevisiae* Spo7p, which is most likely due to the relatively low sequence identity between NEP1R1, Spo7, and other homologs^13^.

The observed interactions between NEP1R1 and CTDNEP1 were largely similar to those predicted by AlphaFold Multimer **(Fig. 6D-E)**, but Alphafold predicted conserved regions 2 (amino acids 101-116) and 3 (amino acids 130-138) of NEP1R1 to fold into an additional helix and a distal beta hairpin **(Supplementary Fig. 5)**. In contrast there was no observable electron density for residues in conserved regions 2 and 3 in the apo CTDNEP1-sNEP1R1 structure, indicating these regions were disordered. However, in the magnesium bound structure, conserved region 2 formed an alpha helix, but differed in its orientation and did not form any significant interactions with CTDNEP1 as predicted by Alphafold **(Supplementary Fig. 5)**.

To assess the functional importance of these interactions, we designed a point mutant, MBP-CTDNEP1ΔAHS232D, to disrupt complex formation. As predicted, MBP-CTDNEP1ΔAHS232D failed to form a stable complex with NEP1R1 on size exclusion chromatography **(Fig. 6F, Supplementary Fig. 1)** and was not activated by sNEP1R1 **(Fig. 6G)**. Consistently, we also recently demonstrated the point mutants F30E in NEP1R1 and V233E in CTDNEP1, which reside at the center of the complex interface, disrupt complex formation in vitro and in mammalian cells with corresponding effects on protein stability and lipin dephosphorylation^16^.

Given that NEP1R1 engages CTDNEP1 at a site far away from the active site and the lack of an experimental CTDNEP1 structure without NEP1R1, it remains difficult to discriminate if NEP1R1 binding activates CTDNEP1 through localized conformational changes and/or by a general global stabilization of the catalytic domain. To assess if NEP1R1 directly stabilized CTDNEP1, we determined the melting temperature (T_m_) of MBP-CTDNEP1ΔAH in the presence and absence of sNEP1R1. We observed that sNEP1R1 increases the thermal stability of MBP-CTDNEP1ΔAH to the same level as the co-purified complex **(Fig. 6H)**. In control experiments, sNEP1R1 did not affect the thermal stability of the MBP-CTDNEP1ΔAH S232D and V233E^16^ point mutants, which disrupt complex formation **(Fig. 6H)**. While we are currently unable to directly assess the specific conformational changes in CTDNEP1 induced by NEP1R1 binding, this confirms that NEP1R1 binding contributes to a global allosteric stabilization and activation of the CTDNEP1 catalytic domain, which is consistent with our previous observations that NEP1R1 prevents CTDNEP1 aggregation.

### Peptide recognition by CTDNEP1

The linker peptide included in the CTDNEP1-sNEP1R1 fusion was involved in a crystal contact and occupied the active site of an adjacent CTDNEP1 molecule in the crystal lattice **(Fig. 7A)**. The linker peptide in the CTDNEP1 active site adopted a similar position as a co-crystallized phospho-peptide substrate complexed with Scp1^32^, which represents a related CTD phosphatase **(Supplementary Fig. 4)**. This suggested the bound linker peptide was serving as a pseudo-substrate and had locked the CTDNEP1-NEP1R1 complex in a catalytically competent position.

**Fig. 7.**
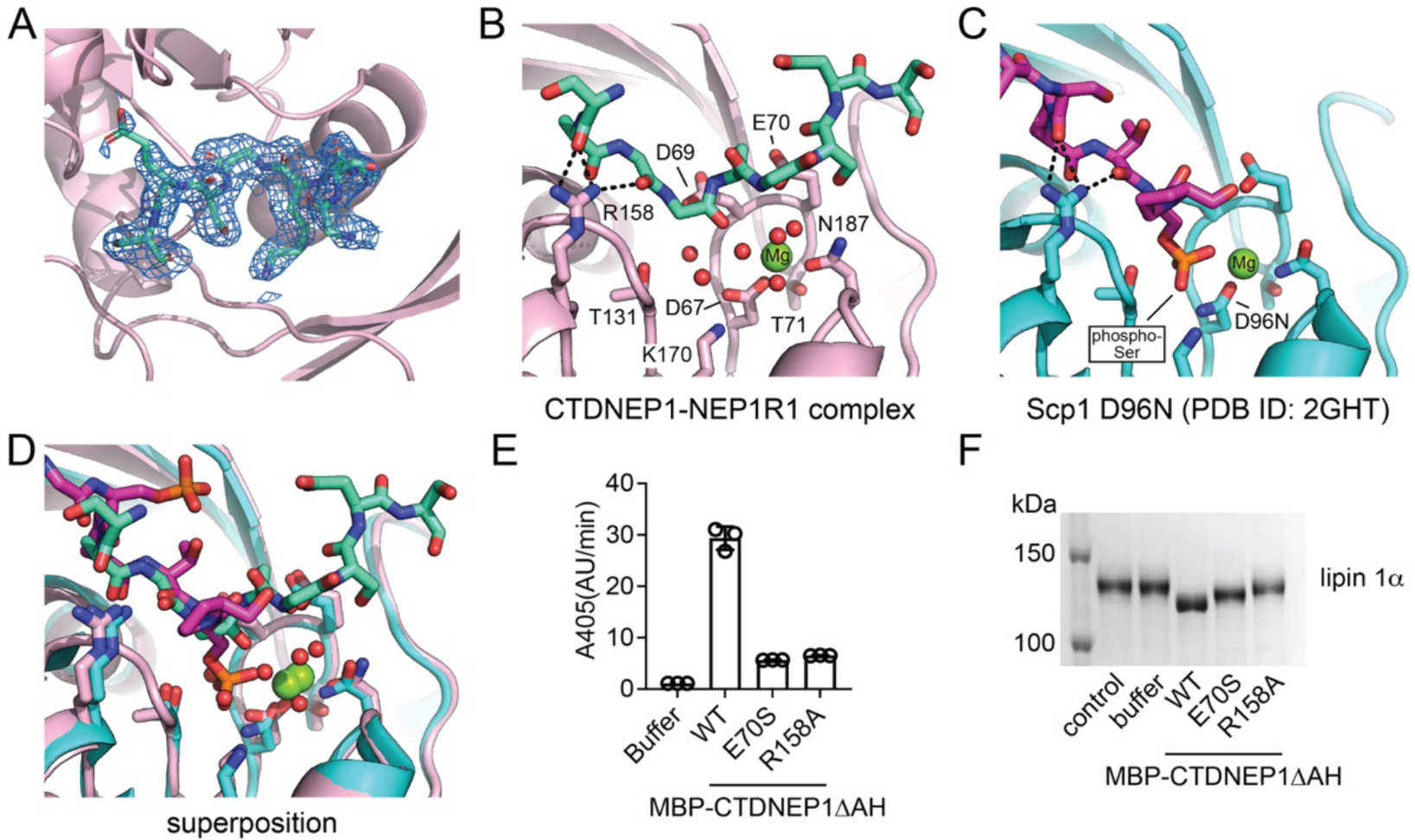
Peptide recognition by CTDNEP1. **(A)** Model of the linker peptide bound in the CTDNEP1 active site with the 2Fo-2Fc electron density map contoured at 1σ. **(B)** CTDNEP1 residues involved in binding the linker peptide. Intermolecular hydrogen bonds between the conserved Arginine residue and the carbonyl groups of the linker peptide are shown as black dotted lines. **(C)** Scp1 residues involved in binding a phospho-peptide. Intermolecular hydrogen bonds between the conserved Arginine residue and the carbonyl groups of the phosphor-peptide are shown as black dotted lines. **(D)** Superimposition of the CTDNEP1 and Scp1 structures with bound peptides. **(E)** Quantification of pNPP hydrolysis by MBP-CTDNEP1ΔAH WT, E70S and R158A. Error bars represent standard deviation (*n* = 3). **(F)** Phos-tag SDS-PAGE analysis of dephosphorylation of mouse lipin 1α after treatment with purified MBP-CTDNEP1ΔAH WT, E70S, and R158A proteins.

In both the CTDNEP1-NEP1R1 and Scp1 structures, a series of intermolecular hydrogen bonds between a conserved arginine residue and carbonyl oxygens of the bound peptides stabilized a single helical turn within the bound peptides that was N-terminal (residues in position −2 to −3) to the site of dephosphorylation **(Fig. 7B, C, D)**. This commonality suggested the conserved arginine residue (R158 in CTDNEP1) was involved in peptide/substrate recognition. To test this hypothesis, we generated the point mutant R158A in MBP-CTDNEP1ΔAH **(Supplementary Fig. 6A)** and assessed activity against pNPP and lipin 1α. To discriminate between general effects on catalysis versus specific effects on substrate recognition, we also generated an E70S point mutant that replaced a conserved glutamate residue in the active site that coordinates a Mg^2+^ bound water molecule and would likely be important for general catalysis, but not peptide binding.

Both the R158A and E70S point mutants had diminished catalytic function, with both point mutants retaining ∼20 % activity compared to WT MBP-CTDNEP1ΔAH using the soluble substrate pNPP **(Fig. 7E)**. In contrast, the R158A point mutant nearly eliminated the ability of MBP-CTDNEP1ΔAH to dephosphorylate lipin 1α **(Fig. 7F)**, while the E70S point mutant only diminished the ability to dephosphorylate lipin 1α, similar to the diminished activity against pNPP. Together, this suggests R158 has two roles in CTDNEP1 catalysis, one to bind and properly orient peptide substrates in the active site for dephosphorylation, and a second involving general stabilizing interactions within CTDNEP1 **(Supplementary Fig. 6B)** that contribute to CTDNEP1 catalytic function.

## DISCUSSION

Prior studies have demonstrated that NEP1R1 acts as a binding partner with CTDNEP1 to promote lipin dephosphorylation and that loss of CTDNEP1, and the consequent inability to dephosphorylate lipin, leads to ER expansion. However, the effects of NEP1R1 depletion in human cells had not been determined, which left open the question if CTDNEP1 requires NEP1R1 in this context. Our finding that depleting NEP1R1 results in ER expansion corroborates that the function of CTDNEP1 and NEP1R1 in regulating ER/NE membrane morphology is evolutionarily conserved with the Nem1-Spo7 complex.

Our biochemical evidence establishes NEP1R1 as an activating regulatory subunit for CTDNEP1, with NEP1R1 directly binding, stabilizing, and enhancing the catalytic activity of CTDNEP1. It is still unclear if the mechanistic role(s) of NEP1R1 also includes to recruit protein substrates for dephosphorylation by CTDNEP1, as seen in some regulatory subunits for canonical phosphoprotein phosphatases ^33,34^. Since NEP1R1 enhances CTDNEP1 mediated catalysis, this hypothesis may be difficult to test without observing a direct protein-protein interaction between NEP1R1 and a CTDNEP1 substrate (e.g. lipin). We did observe that recombinant NEP1R1 was more stable in vitro than CTDNEP1, which required fusion with MBP to prevent aggregation and/or precipitation. Complex formation with NEP1R1 directly stabilized CTDNEP1 and alleviated the propensity of CTDNEP1 to aggregate. This is consistent with recent findings that NEP1R1 regulates the stability and degradation of CTDNEP1 in human cells^16^. Thus, we suspect that NEP1R1 may have several pleiotropic roles that contribute to CTDNEP1 cellular function, in addition to directly increasing CTDNEP1 catalysis.

While NEP1R1’s cellular concentration has not been determined, CTDNEP1’s cellular concentration in HEK293 cells is estimated to be 19 nM^35^, which is two orders of magnitude lower than the K_d_ of ∼2.9 μM between CTDNEP1ΔAH and sNEP1R1. This raises the likely possibility that CTDNEP1 can dissociate from NEP1R1 in cells, which has recently been demonstrated^16^. Dissociation from NEP1R1 may be a critical factor in regulating CTDNEP1 function, which could either lead to CTDNEP1 protein degradation^16^, independent functions of CTDNEP1 (e.g dephosphorylation of nuclear myc^20^), or association with other yet identified regulatory subunits. Future studies addressing these issues may take advantage of the soluble complex we identified that is sufficient for complex formation, and/or the fusion complex of CTDNEP1-NEP1R1 that covalently links the two proteins together and increases thermal stability.

While the K_d_ of the orthologous Nem1-Spo7 complex has not been determined, it has been suggested that Nem1-Spo7 form a constitutive complex. In yeast, Ice2 has been identified as a regulator of Nem1-Spo7 that inhibits function^36^. The mammalian SERINC proteins display sequence and structural homology with Ice2^37^, however it is not known if they analogously inhibit CTDNEP1-NEP1R1, in addition to their reported role of incorporating serine into membrane lipid synthesis^38^. However, dissociation of CTDNEP1 and NEP1R1 suggests that this need not necessarily be the case, as CTDNEP1-NEP1R1 may have evolved different regulatory mechanisms from their yeast counterparts.

Comparison of the experimentally determined crystal structure versus the AlphaFold Multimer prediction identified important shared features for complex formation, but also had some notable differences. Alphafold predicted NEP1R1 to form a cytoplasmic domain when bound to CTDNEP1, whereas our experimental structure lacked density beyond the single helix formed by conserved region 1 or had a second helix that adopted a different orientation to that predicted by AlphaFold. Mutational analyses confirmed that the shared residue interactions found in both the experimental and predicted complexes are necessary for complex formation, but the discrepancies between the experimental and AI structural predictions remain unresolved. Additional experimental structures of either the full-length NEP1R1-CTDNEP1 complex or a non-fused complex may resolve these issues.

Some phosphoprotein phosphatases bind their regulatory subunits using a consensus motif^39,40^. At this point, given the lack of other known CTDNEP1 regulatory subunits, it is not possible to predict a consensus motif for CTDNEP1 regulatory subunits. However, based on the CTDNEP1-NEP1R1 binding interface it remains possible that other proteins may associate with CTDNEP1 through a helix that would bind and occlude the hydrophobic patch on CTDNEP1 to mediate complex formation.

Lastly, inactivating truncations and point mutations in CTDNEP1 have recently been identified in patients with medulloblastoma, an aggressive brain cancer. Mapping the location of these point mutants provides a clear rationale for loss of function. Both point mutants, L72H and W205R are in the core of the catalytic domain and would likely disrupt correct protein folding and adoption of the native tertiary structure necessary for catalysis **(Supplementary Fig. 7)**. Notably, both mutations are located at sites away from the NEP1R1 binding interface, and mutations of NEP1R1 in medulloblastoma have not been identified. Thus, it is unclear if NEP1R1 is required for CTDNEP1 to function as a tumor suppressor in brain cancer cells, if another regulatory subunit is necessary, or if CTDNEP1 alone is sufficient.

## METHODS

### Materials

Lambda Protein Phosphatase, Amylose resin, Q5 Site-Directed Mutagenesis Kit and NEBuilder® HiFi DNA Assembly Master Mix were purchased from New England Biolabs. 4-Nitrophenyl phosphate (pNPP), Nickel-NTA Agarose resin and n-Dodecyl-β-D-Maltopyranoside (DDM) were from Gold Biotechnology. n-Decyl-β-D-Maltopyranoside (DM), and glyco-diosgenin (GDN) were products of Anatrace. POPA (1-palmitoyl-2-oleoyl-sn-glycero-3-phosphate, #840857), POPE (1-palmitoyl-2-oleoyl-sn-glycero-3-phosphoethanolamine, #850757) and POPC (1-palmitoyl-2-oleoyl-glycero-3-phosphocholine, #850457) were from Avanti Polar Lipids. SYPRO orange protein gel stain was from Invitrogen.

### Plasmids

Plasmids were constructed using ligation independent cloning (LIC), site-directed mutagenesis, or Gibson assembly. pET-His6-MBP-TEV LIC cloning vector (1M) was a gift from Scott Gradia (Addgene plasmid #29656; http://n2t.net/addgene:29656; RRID: Addgene_29656). pET-His6 msfGFP-TEV cloning vector with BioBrick polycistronic restriction sites (9GFP) was a gift from Scott Gradia (Addgene plasmid #48287; http://n2t.net/addgene:48287; RRID: Addgene_48287).

The following is the list of plasmids used in this research.

His6-SUMO-NEP1R1
His6-MBP-TEV-CTDNEP1
His6-MBP-TEV-CTDNEP1ΔAH (delete 1-39 amino acids from CTDNEP1)
His6-MBP-TEV-CTDNEP1ΔAHE70S
His6-MBP-TEV-CTDNEP1ΔAHR158A
His6-MBP-TEV-CTDNEP1ΔAHS232D
His6-MBP-TEV-NEP1R1/CTDNEP1 coexpression
His6-MBP-TEV-NEP1R1/CTDNEP1D67E coexpression
His6-MBP-TEV-NEP1R1/CTDNEP1ΔAH coexpression
His6-msfGFP-MBP-TEV-CTDNEP1ΔAH
His6-TEV-sNEP1R1 (delete 9-25 and 43-107 amino acids)
His6-MBP-TEV-sNEP1R1/CTDNEP1ΔAH coexpression
His6-sNEP1R1/CTDNEP1ΔAH coexpression
CTDNEP1ΔAH-linker-sNEP1R1-His6 (linker amino acid sequence: GSAKGSES)

### Mammalian cell lines

U2 OS and HEK293-T cells were grown at 37 °C in 5 % CO_2_ in DMEM low glucose (Gibco 11885) supplemented with 10 % heat inactivated FBS (F4135) and 1 % antibiotic-antimycotic (Gibco 15240112). Cells were cultured without antibiotics during RNAi treatments for experiments. Cells were used for experiments before passage 25. Cells were tested for mycoplasma upon initial thaw and generation of new cell lines (Southern Biotech 13100-01), and untreated cells were continuously profiled for contamination by assessment of extranuclear DAPI/Hoechst 33258 staining.

### Stable cell line generation

U2OS CTDNEP1^KO^ + CTDNEP1-HA + Flag-NEP1R1 stable cells were generated by retroviral transduction, and bulk populations of cells were used for experiments. Retroviruses were generated by transfecting HEK293T cells with pCG-gag-pol, pCG-VSVG and pMRX-Flag-NEP1R1-Neo using Lipofectamine 2000. The retroviruses were recovered 48hrs post-transfection, filtered using a 0.22 μm PVDF syringe filter and used to transduce U2OS CTDNEP1^KO^ + CTDNEP1-HA stable cells. After 48 h of infection, cells were placed under 300 μg/mL G418 selection for ∼1 week or until control cells where dead, then frozen and/or used for experiments. Cells were continuously cultured in 7.5 μg/mL blasticidin + 300 μg/mL G418.

### Immunoblot

Lysis buffers used: 0.1 % Triton X-100, 50 mM NaF, 1 mM EDTA, 1 mM EGTA, 10 mM Na_2_HPO_4_, 50 mM β-glycerophosphate, 1 tablet/50 mL cOmplete protease inhibitor cocktail (Roche), pH 7.4. Cell lysates were obtained by adding lysis buffer to cell pellets collected by trypsinization and centrifugation at 300 x g for 5 min followed by 1-2 PBS washes. Lysates were homogenized by pushing through a 23G needle 30 times and then centrifuged at >20,000 x g for 10 min at 4 °C, then protein concentration was determined using the Pierce BCA Protein assay kit (Thermo Scientific 23225). 20-30 mg of whole cell lysates/lane were run on 8-15% polyacrylamide gels dependent on target size, and protein was wet transferred to 0.22 mm nitrocellulose. Ponceau S staining was used to visualize transfer efficiency, then washed with TBS or DI water; then, membranes were blocked in 5 % nonfat dry milk or BSA in TBS for 1 h. Membranes were then incubated with primary antibodies in 5 % milk or BSA for 1-2 hours at room temperature. Membranes were washed 3 times for 5 min in TBS-T, then incubated with anti-HRP secondary antibodies in 5 % milk or BSA in TBS-T for 1 h at room temperature with rocking. Membranes were washed 3 times for 5 min in TBS-T. Clarity or Clarity Max ECL reagent (Bio-Rad 1705060S, 1705062S) was used to visualize chemiluminescence, and images were taken with a Bio-Rad ChemiDoc or ChemiDoc XRS+ system. Exposure times of images used for analysis or presentation were maximum exposure before saturation of pixels around or within target bands. Antibody concentrations used: Mouse anti-a tubulin DM1A 1:5000; Mouse anti-Flag 1:4000; Rabbit anti-HA 1:1000; all secondaries 1:10000.

### Immunofluorescence

Cells were washed 2x with warm PBS and fixed in 4 % paraformaldehyde + 0.1 % glutaraldehyde in PBS for 15 min, permeabilized in 0.5 % Triton X-100 for 5 min, then washed 3 times with PBS and blocked in 2 % BSA in PBS for 30 min. Samples were transferred to a humidity chamber and incubated with primary antibodies in 2 % BSA in PBS for 1 h at room temperature with rocking. Samples were washed with PBS 3 times for 5 min, then incubated with secondary antibodies in 2 % BSA in PBS for 1 h at room temperature in the dark with rocking. Samples were then washed with PBS 3 times for 5 min in the dark. Coverslips were mounted with ProLong Gold Antifade reagent + DAPI (Thermo Fisher P36935) and sealed with clear nail polish.

### Microscopy

Samples were imaged on an inverted Nikon Ti microscope equipped with a Yokogawa CSU-X1 confocal scanner unit with solid state 100-mW 488-nm and 50-mW 561-nm lasers, using a 60×1.4 NA plan Apo objective lens (or 10x 0.25 NA ADL objective with 1.5x magnification), and a Hamamatsu ORCA R-2 Digital CCD Camera.

### Image analysis

Image analysis was performed using FIJI/ImageJ. For all scoring phenotypes quantified by categorization were scored blindly. Images were blinded for analysis using the ImageJ Macro ImageJ Filename_Randomizer, cells where randomized and analysis was done blindly. GraphPad Prism 8 was used for all statistical analysis. In imaging experiments where phenotypes of individual cells are scored, n refers to individual cells. All N refer to experimental repeats. p values, Fisher’s exact tests for ER expansion phenotype.

### Protein expression

Plasmids were transformed into BL21 (DE3) RIPL cells and inoculated into Terrific Broth containing 50 μg/mL kanamycin and cultured overnight at 37 °C. 10 mL overnight culture was transferred to 1 L Terrific Broth with 50 μg/mL kanamycin, cultured at 37 °C to an OD600 between 1.8 and 2.2, then switch the culture temperature to 15 °C till the culture temperature reach 20 °C, then isopropyl β-D-1-thiogalactopyranoside (IPTG) was added to a final concentration of 0.5 mM, keep the culture temperature at 20 °C for 18 to 22 h. Cells were collected by centrifugation at 4,000 x g for 20 min, cells pellets were stored at −80 °C.

### Protein Purification

All the purification steps were carried out at 4 °C or on ice, unless otherwise specified. The following buffers were used. Buffer A: 20 mM Tris pH 7.5, 150 mM NaCl, 10 % glycerol, and 10 mM beta-mercaptoethanol (BME); Buffer B: 20 mM Tris pH 7.5, 300 mM NaCl, 20 mM imidazole, and 10 mM BME; Buffer C: 20 mM Tris pH 7.5, 100 mM NaCl, 300 mM imidazole, and 10 mM BME; and Buffer D: 20 mM Tris pH 7.5, 100 mM NaCl, and 10 mM BME.

His6-MBP-NEP1R1/CTDNEP1 co-expression, His6-MBP-NEP1R1/CTDNEP1ΔAH co-expression, His6-MBP-CTDNEP1 and His6-SUMO-NEP1R1 were purified as following. Cell pellets were lysed in buffer A by sonication, and lysates were centrifuged at 10,000 x g for 30 min. Supernatant was collected and centrifuged at 100,000 x g for 1 h. The pellet was resuspended in buffer A containing 1 % DDM (n-Dodecyl-beta-maltoside) and extracted at 4 °C by rotating overnight, 20 mM imidazole was added to the extraction before spining at 100,000 x g for 40 min, supernatant was collected and incubated with Ni-NTA resin for 1 h, the resin was then transferred to an empty gravity column, washed with buffer B containing 0.05 % DDM, then the protein was eluted with buffer C containing 0.05 % DDM. His-SUMO-NEP1R1 elution from Ni-NTA was digested with ULP1 (Ubl-specific protease 1) at 4 °C overnight, then loaded onto Superdex 200 16/600 HiLoad (Cytiva), and eluted with buffer D containing 0.05 % DDM. Target proteins with MBP-tag were loaded onto amylose resin after elution from Ni-NTA, washed with buffer D containing 0.05 % DDM, and eluted with 10 mM maltose in buffer D supplied with 0.05 % DDM. The eluted protein was further purified by size exclusion chromatography on Superdex 200 16/600 HiLoad equilibrated with buffer D containing 0.02 % glyco-diosgenin (GDN). Fractions containing the target proteins were collected, concentrated and flash-frozen in liquid nitrogen, then stored at −80 °C.

All other proteins were purified using the following protocol. Cell pellets were lysed in buffer A by sonication, then imidazole were added into the lysates to final concentration of 20 mM before centrifugation at 100,000 x g for 30 min. Supernatant was collected and incubated with Ni-NTA resin for 1 h, the resin was then transferred to an empty gravity column, washed with buffer B, then eluted with buffer C. Proteins with MBP-tag were loaded onto amylose resin (New England Biolabs) after elution from Ni-NTA, washed with buffer D, and eluted with buffer D supplied with 10 mM maltose. The eluted protein was further purified by size exclusion chromatography on Superdex 75 26/600 HiLoad (Cytiva) or Superdex 200 26/600 HiLoad (Cytiva) equilibrated with buffer D. Fractions containing the target proteins were collected, concentrated and flash-frozen in liquid nitrogen, then stored at −80 °C.

### Lipin purification

Full-length mouse lipin 1α was purified as recently described^41^. Briefly, lipin 1α was expressed in Sf9 cells using baculovirus. 300 mL of cells were infected with 1.0 mL of baculovirus at 3 million cells/mL at >95% viability and harvested 72 h later. Cell pellets were lysed in 40 mM Tris, pH 8.0, 300 mM NaCl, 10 mM BME by sonication, and the lysates were centrifuged at 81,770 × g for 30 min. Lipin 1α protein was purified using Ni-NTA resin. Eluted protein was applied to Streptactin-XT resin (IBA lifesciences) equilibrated with equilibration buffer (100 mM Tris pH 8.0, 150 mM NaCl, 10 mM BME), then washed with equilibration buffer, and eluted with equilibration buffer containing 50 mM biotin. Proteins were further purified by size exclusion chromatography using a Superdex 200 increase 10/300 GL column (Cytiva) in 20 mM Tris pH 8.0, 150 mM NaCl, and 10 mM BME. Fractions containing lipin 1α protein were concentrated, flash-frozen, and stored at −80 °C.

### pNPP assay

100 mM pNPP was prepared in 50 mM HEPES pH 6.7, 100 mM NaCl, 10 mM MgCl_2_, 10 mM BME. The reactions were carried out in 96-well plate by mixing 95 μL pNPP solution with 5 μL enzyme (0.1 μM final protein concentration), and the absorbance at 405 nm was monitored with SpectraMax M2e Microplate Readers (Molecular Devices) at 30 sec intervals at ambient temperature for 30 min.

### Interaction of His6-MBP-CTDNEP1ΔAH with NEP1R1

Purified His6-MBP-CTDNEP1ΔAH or His6-MBP-CTDNEP1ΔAHS232D (12 μM, final protein concentration) were mixed with NEP1R1 (53 μM, final protein concentration), incubated on ice for 3 h, and the mixture was centrifuged at 15,000 rpm for 15 min at 4 °C. The supernatant was loaded onto a Superdex200 increase 10/300 size-exclusion chromatography column. Phosphatase activity of each fraction was analyzed using the pNPP assay, and protein composition of each fraction was resolved by SDS-PAGE

### Microscale thermophoresis

Microscale thermophoresis^42^ (MST) experiments were carried out on a Monolith NT.115 (NanoTemper Technologies) using standard capillaries with the following settings: excitation: Nano-blue at 5% LED power and medium MST power. The buffer consisted of 20 mM Tris pH 7.5, 100 mM NaCl, 10mM BME for the assay. Each capillary contained His6-msfGFP-MBP-CTDNEP1ΔAH at 1.0 μM that was mixed with an equal volume of sNEP1R1. The highest concentration of sNEP1R1 was 0.4 mM and lower concentrations were generated by 16 x 1:1 serial dilutions. The final concentration of His6-msfGFP-MBP-CTDNEP1ΔAH was 0.5 μM and the concentration of sNEP1R1 was varied between 6.1 nM and 0.2 mM. The mixtures were incubated for 15 min at room temperature and the measurement was conducted at ambient temperature. The data were analyzed with MO.Affinity Analysis software version 2.3 (NanoTemper Technologies) using the signal from an MST-on time of 1.5 s. Four independent experiments were conducted. Error bars represent standard deviation.

### Lipin dephosphorylation

Purified mouse lipin1α (5 μM) was incubated with 5 μM of enzyme in 20 mM Tris pH 7.5, 100 mM NaCl, 10 mM MgCl_2_, 10 mM BME at 30 °C for 30 min. Lipin (5 μM) treated with lamda protein phosphatase (800 U) was carried out according to the product protocol, incubated at 30 °C for 30 min. An equal volume of 2X SDS-sample buffer were added to the reaction. The dephosphorylation of lipin was analyzed on phos-Tag SDS-PAGE.

### Liposome sedimentation assay

Liposomes were prepared using a thin film hydration method. Briefly, POPC, POPE and POPA in chloroform were mixed at the desired molar ratios (liposome composed of POPC and POPE with a molar ratio of 8:2, while liposome containing POPC, POPE and POPA with a molar ratio of 7:2:1) and then dried under nitrogen gas. The dried lipids were resuspended in a buffer containing 50 mM Tris pH 7.5, 100 mM NaCl, 10 mM BME. Large unilamellar vesicles (LUVs) were generated by seven freeze-thaw cycles, then sonicated in a water bath for 10 min. Liposome and protein were mixed to give a final concentration of 1.0 mM liposomes and 1.0 µM protein. The mixture was incubated for 30 min at 4 °C and centrifuged at 100,000 × g at 4 °C for 1 h using a TLA100 fixed angle rotor (Beckman). The supernatant fraction was carefully removed, and the protein content of the pellet and supernatant fractions were analyzed by SDS-PAGE. All binding assays were performed at least three times and SDS-PAGE gel bands were quantified using ImageJ.

### Thermal shift assay

MBP, MBP-CTDNEP1ΔAH, MBP-CTDNEP1ΔAH S232D and MBP-CTDNEP1ΔAH V233E were incubated with His-TEV-sNEP1R1 at 1:1 molar ratio (final concentration was 10 μM of each) at 4 °C for 1 h, then equal volume of diluted SYPRO orange dye were mixed with the protein solution. The Thermal shift assay was carried out with StepOneplus Realtime PCR system (Thermo fischer Scientific) under the following settings: Temperature range: 25-70 °C with a temperature increase of 0.5 °C/min. The melting temperature (Tm) was analyzed with StepOne software.

### Crystallization and data collection

Purified CTDNEP1ΔAH-sNEP1R1-His6 fusion protein was used for crystallization. All crystals were grown using the hanging-drop method by mixing 1.5 μL of reservoir solution with 1.5 μL of protein solution at room temperature. Crystals were grown in 5.5 – 6 % PEG3350, 0.2 M lithium citrate, 0.3 - 0.4 % CHAPS with or without seeding. Crystals of the apo CTDNEP1ΔAH-sNEP1R1-His6 fusion were cryoprotected with 0.1 M lithium citrate, 6 % PEG3350, 30 % PEG400 and flash frozen in liquid nitrogen prior to data collection. Crystals of Mg^2+^ bound to CTDNEP1ΔAH-sNEP1R1-His6 fusion were first soaked in a solution of 40 % PEG400 with 0.5 M MgCl_2_ and cryoprotected with 40 % PEG400 and 0.1 M MgCl_2_. Diffraction data were collected at Brookhaven National Lab NSLS II AMX beamline 17-ID-1 and processed using the AutoProc pipeline^43^.

### Structure determination and refinement

Phases were determined by molecular replacement in Phenix^44^ using Phaser^45^. A truncated CTDNEP1 alphafold^29^ model with B-factors adjustments using Phenix was used as a search model. Additional model building in Coot^46^ and refinement in Phenix produced the final apo model of the CTDNEP1-NEP1R1 fusion (**Table 1**, PDB code: 8UJL). The Mg^2+^ bound structure was phased by molecular replacement with the final apo model as the search model. The final Mg^2+^ bound model was produced by manual model building in coot and refinement in Phenix (**Table 1**, PDB code: 8UJM). The Mg^2+^ ion was modeled based on the strongest positive peak in an Fo−Fc difference map after initial refinement and geometry and distance restraints were used in refinement.

## ACKNOWLEDGEMENTS

We thank Jean Jakoncic and Alexei Soares (Brookhaven National Lab) for their assistance with data collection.

