## Supplementary Info for "Structure and mechanism of the human CTDNEP1-NEP1R1 membrane protein phosphatase complex necessary to maintain ER membrane morphology"

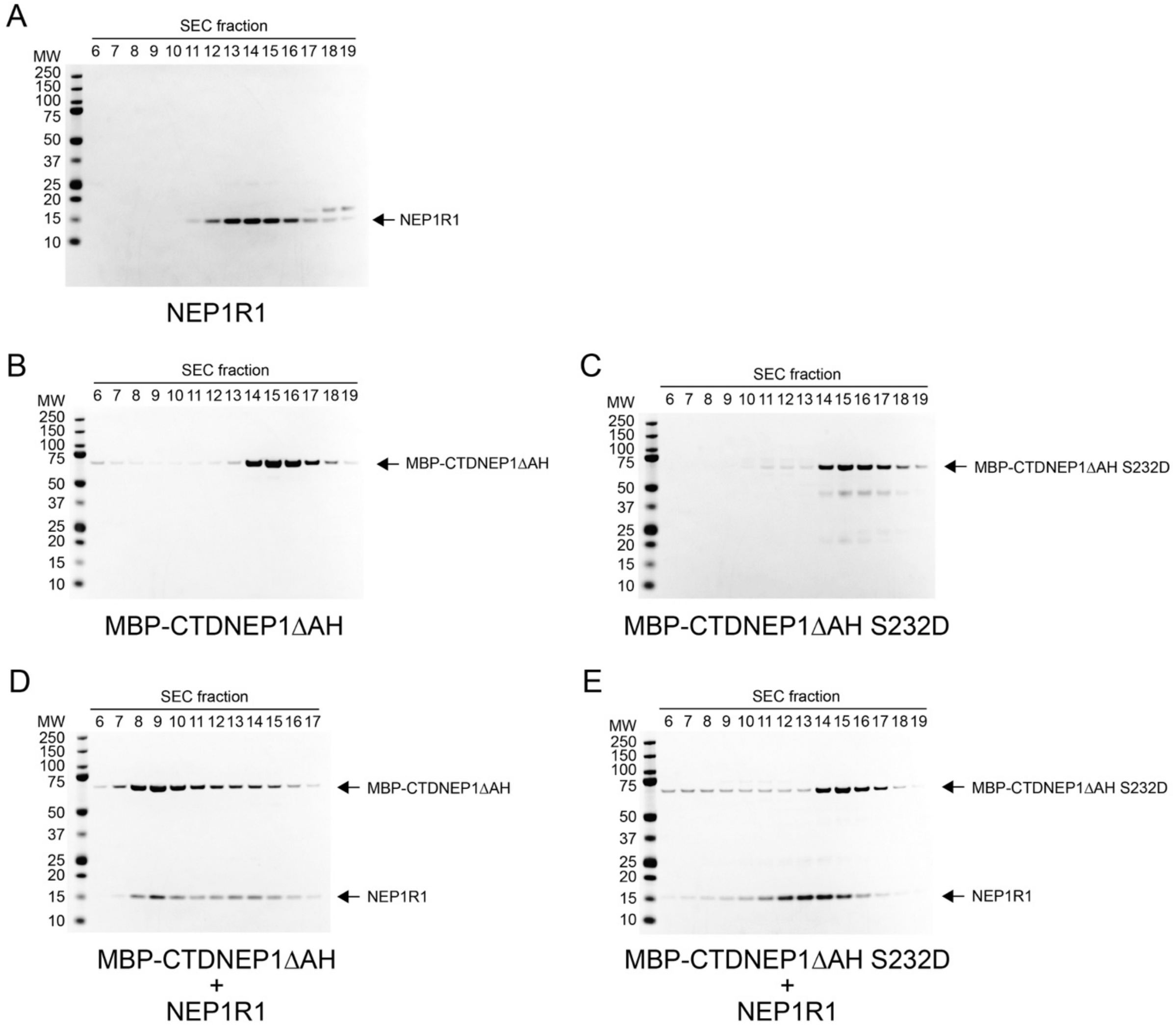


**Supplementary Figure 1. CTDNEP1 and NEP1R1 form a stable complex on size-exclusion chromatography that can be disrupted by mutation of the binding interface.** SDS-PAGE analysis of CTDNEP1-NEP1R1 complex formation using fractions from size-exclusion chromatography of **(A)** NEP1R1 alone, **(B)** MBP-CTDNEP1ΔAH alone, **(C)** MBP-CTDNEP1ΔAH S232D, **(D)** a mixture of MBP-CTDNEP1ΔAH and NEP1R1 in a 1:4 molar ratio, and **(E)** a mixture of MBP-CTDNEP1ΔAH S232D and NEP1R1 in a 1:4 molar ratio.

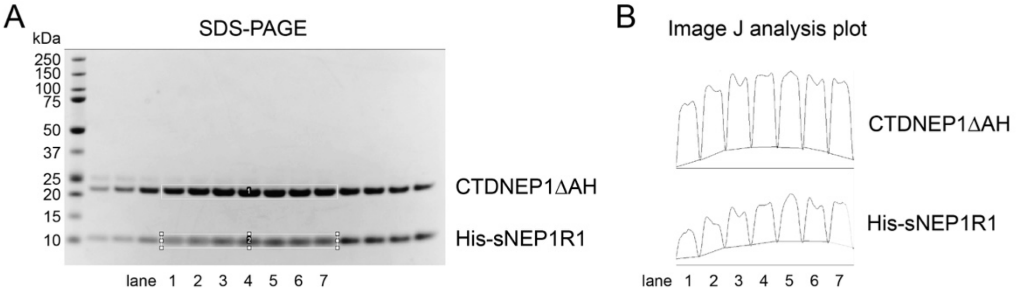


C

Image J analysis results

| lane | CTDNEP1ΔAH (peak integration) | His-sNEP1R1 (peak integration) | Experimental ratio CTDNEP1ΔAH / His-sNEP1R1 |
|---|---|---|---|
| 1 | 14035 | 5499 | 2.55 |
| 2 | 16243 | 7094 | 2.29 |
| 3 | 17779 | 8378 | 2.12 |
| 4 | 18486 | 9144 | 2.02 |
| 5 | 17923 | 9520 | 1.88 |
| 6 | 17531 | 8858 | 1.98 |
| 7 | 17995 | 9375 | 1.92 |

| | |
|---|---|
| Experimental ratio (average) | 2.11 |

D

| | MW (Daltons) |
|---|---|
| CTDNEP1ΔAH | 23742 |
| His-sNEP1R1 | 10240 |

| | |
|---|---|
| Theoretical ratio for 1:1 complex | 2.32 |

**Supplementary Figure 2. Stoichiometry analysis of subunit composition for the CTDNEP1ΔAH-sNEP1R1 protein phosphatase complex. (A)** SDS-PAGE analysis of the size-exclusion fractions of untagged CTDNEP1ΔAH and His-sNEP1R1 after co-expression and purification using Ni-NTA. **(B)** Image J analysis plot of the SDS-PAGE band intensities for CTDNEP1ΔAH and His-sNEP1R1 from the indicated lanes in panel A. **(C)** Quantitation of the Image J analysis indicates CTDNEP1ΔAH and His-sNEP1R1 form a complex in a 1:1 molar ratio with the experimental ratio of band intensities similar to **(D)** the theoretical ratio of band intensities for a 1:1 complex of CTDNEP1ΔAH (MW=23.742 kDa) and His-sNEP1R1 (MW=10.24 kDa) based on their relative molecular weights.

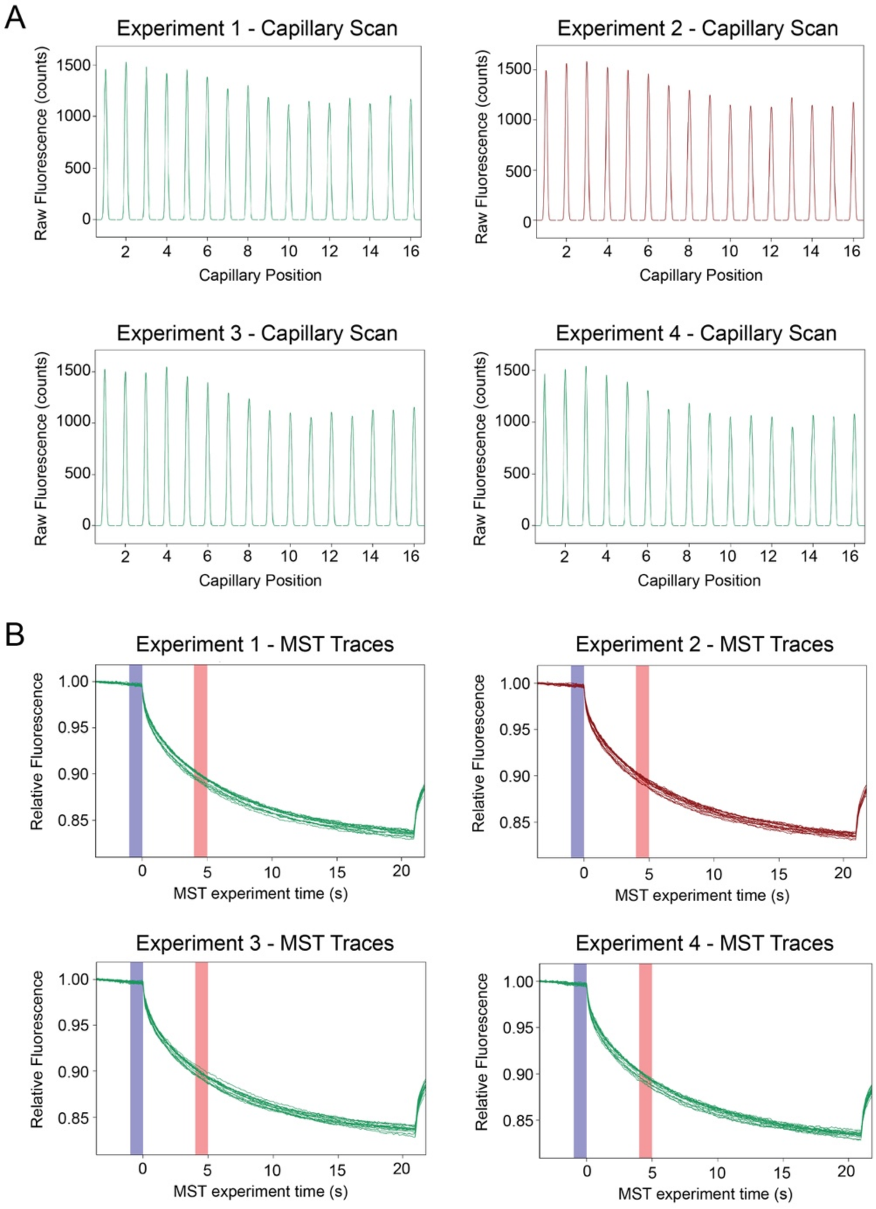


**Supplementary Figure 3. Microscale thermophoresis (MST). (A)** Raw fluorescence readings for individual capillaries containing a constant concentration of 0.5 μM msfGFP-MBP-CTDNEP1ΔAH and increasing concentrations of His-sNEP1R1 that varied between 6.1 nM and 0.2 mM. The four panels represent individual experimental replicates (*n* = 4). **(B)** Plots (*n* = 4) of the MST time traces (green or red lines) on one graph for the individual capillaries with increasing concentrations of His-sNEP1R1. The measured fluorescence from the msfGFP-MBP-CTDNEP1ΔAH molecule changes based on the movement profile in a temperature gradient as a function of size and complex formation with His-sNEP1R1. The vertical blue band indicates the IR-laser was off ($F_{cold}$), prior to turning the IR-laser on. The vertical red band indicates MST-on time of 1.5 sec ($F_{hot}$). The IR-laser was turned off after ~20 sec and the molecules diffused back.

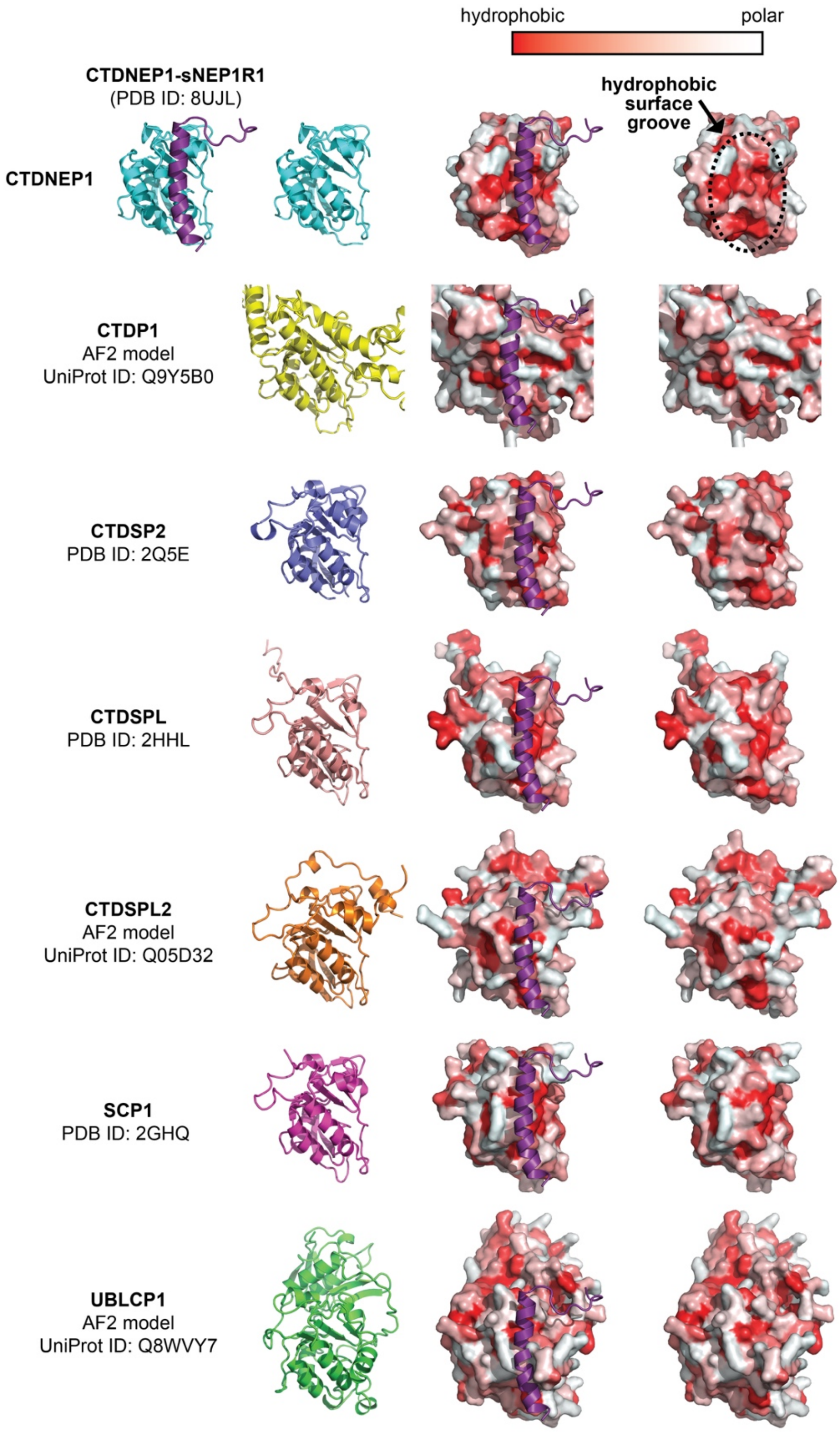


**Supplementary Figure 4. CTDNEP1 associates with NEP1R1 through a hydrophobic surface groove that is not conserved in the CTD phosphatase family.** NEP1R1 (purple) binds to a hydrophobic surface groove on CTDNEP1 (top row). This hydrophobic surface is not conserved in the other 6 members of the CTD phosphatase family (CTDP1, CTDSP2, CTDSPL, CTDSPL2, SCP1, and UBLCP1) that are shown in cartoon (left) and surface views (middle, right) with and without NEP1R1 (purple) superimposed. Hydrophobic residues are red, polar residues are white. AlphaFold2 predictions used when experimental structures were not available.

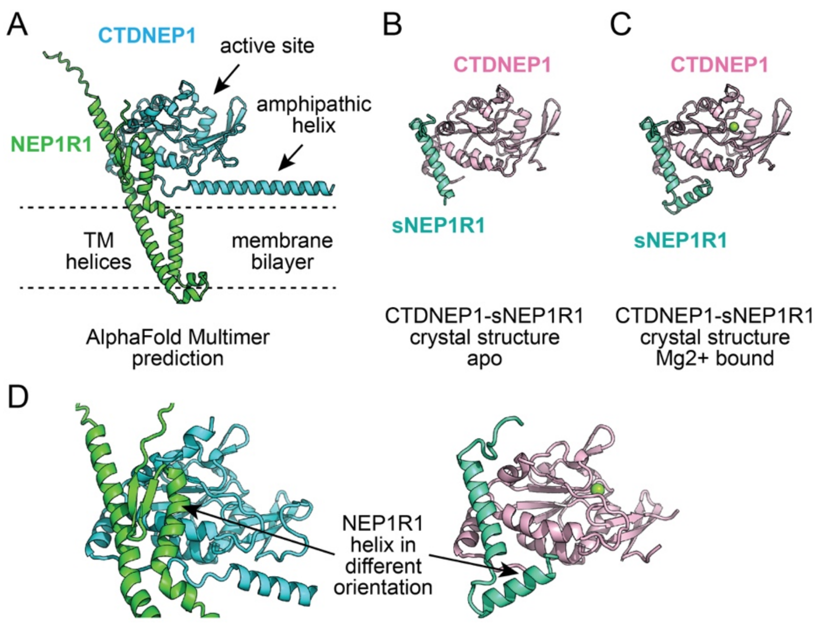


**Supplementary Figure 5. Comparison of CTDNEP1-NEP1R1 experimental crystal structures with an AlphaFold Multimer prediction. (A)** AlphaFold Multimer structural prediction of the CTDNEP1-NEP1R1 complex. The catalytic subunit CTDNEP1 is predicted to adopt a HAD-like phosphatase core with an N-terminal amphipathic helix that would run parallel to the membrane. NEP1R1 is predicted to contain a soluble cytoplasmic domain with two helices and a two-stranded beta sheet, and two transmembrane (TM) helices embedded in the membrane bilayer. **(B, C)** High resolution crystal structures of the CTDNEP1-sNEP1R1 fusion complex **(B)** without and **(C)** with $Mg^{2+}$ bound in the active site. The $Mg^{2+}$ bound structure had electron density for a second helix that did not associate with CTDNEP1. **(D)** Comparison of the AlphaFold multimer prediction (left) with the $Mg^{2+}$ bound crystal structure of CTDNEP1-sNEP1R1 (right) that highlights the different orientation of an NEP1R1 helix predicted to interact with CTDNEP1 by AF Multimer but that was not observed to interact with CTDNEP1 in the experimental crystal structure.

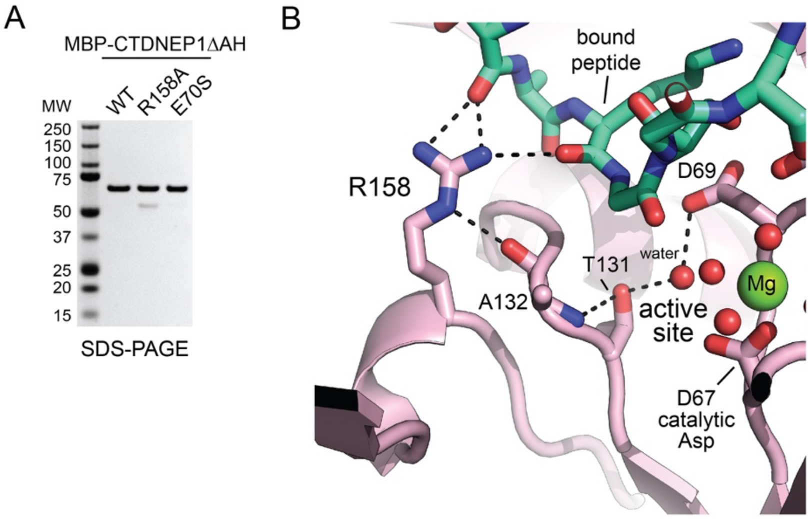


**Supplementary Figure 6. Arg158 has dual structural roles in peptide recognition and stabilizing the active site through an intramolecular hydrogen bond network. (A)** SDS-PAGE analysis of purified MBP-CTDNEP1ΔAH proteins used in this study. **(B)** Zoom in of the interactions of R158 in CTDNEP1 with the bound peptide linker (green) and the main chain carbonyl-oxygen of A132, which helps stabilize the active site through a hydrogen bond network. The point mutant R158A would eliminate all of these interactions to not only affect peptide dephosphorylation, as seen with lipin 1$\alpha$, but also general catalysis towards pNPP.

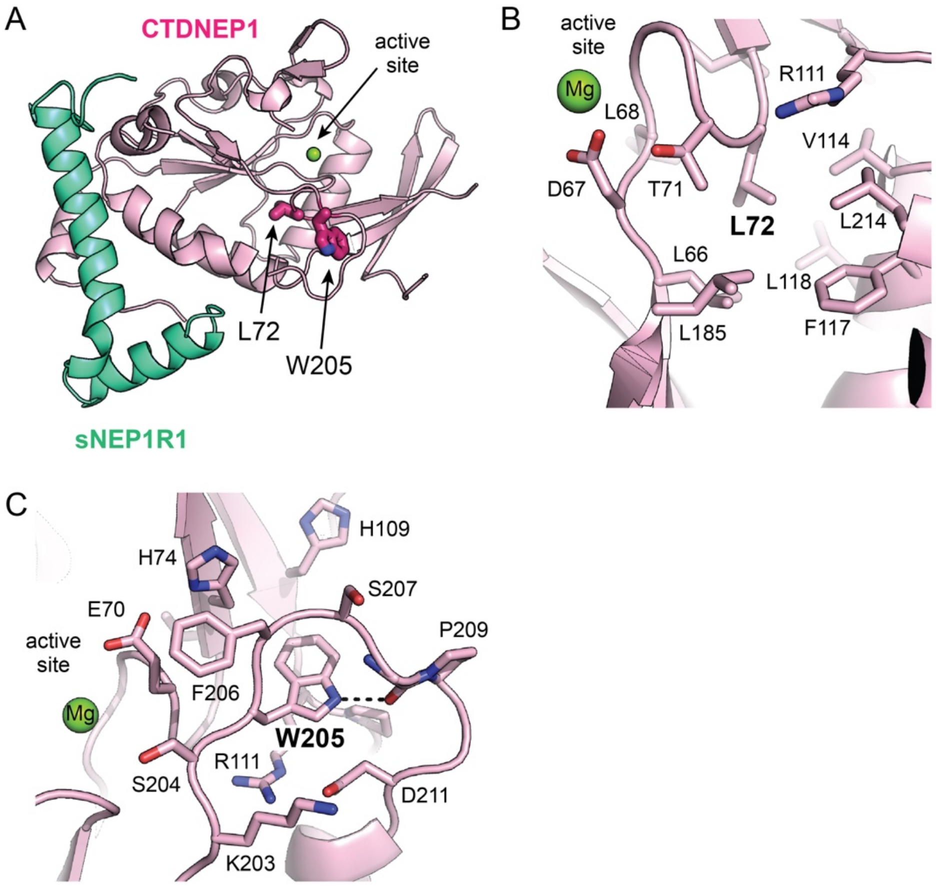


**Supplementary Figure 7. Cancer-associated loss-of-function point mutants in CTDNEP1 are predicted to affect catalysis through destabilization of the protein fold and active site. (A)** Crystal structure of the $Mg^{2+}$ bound CTDNEP1-sNEP1R1 complex that highlights the interior positions of L72 and W205 (magenta sticks), whose mutations are associated with CTDNEP1 loss of function and medulloblastoma. **(B)** Zoom in of the hydrophobic pocket that L72 occupies in the interior of the CTDNEP1 catalytic subunit. The cancer associated mutation L72H would place a polar Histidine residue in the hydrophobic pocket, which is predicted to destabilize the protein fold and affect catalysis by altering the architecture of adjacent active site. **(C)** Zoom in of the interactions of W205. The cancer associated mutation W205R would eliminate the hydrogen bond formed between W205 and a main chain carbonyl-oxygen, and place a positively charged Arg residue in a hydrophobic pocket. This is predicted to disrupt the architecture of the nearby active site and reduce CTDNEP1-mediated dephosphorylation.
